# Pan-cancer landscape of homologous recombination deficiency

**DOI:** 10.1101/2020.01.13.905026

**Authors:** Luan Nguyen, John Martens, Arne Van Hoeck, Edwin Cuppen

## Abstract

Homologous recombination deficiency (HRD) results in impaired double strand break repair and is a frequent driver of tumorigenesis. Here, we developed a genome-wide mutational scar-based pan-cancer **C**lassifier of **HO**mologous **R**ecombination **D**eficiency (CHORD) that can discriminate BRCA1- and BRCA2-subtypes. Analysis of a metastatic (n=3,504) and primary (n=1,854) pan-cancer cohort revealed HRD was most frequent in ovarian and breast cancer, followed by pancreatic and prostate cancer. Biallelic inactivation of BRCA1, BRCA2, RAD51C or PALB2 was the most common genetic cause of HRD, with RAD51C and PALB2 inactivation resulting in BRCA2-type HRD. While the specific genetic cause of HRD was cancer type specific, biallelic inactivation was predominantly associated with loss-of-heterozygosity (LOH), with increased contribution of deep deletions in prostate cancer. Our results demonstrate the value of pan-cancer genomics-based HRD testing and its potential diagnostic value for patient stratification towards treatment with e.g. poly ADP-ribose polymerase inhibitors (PARPi).

## Introduction

The homologous recombination (HR) pathway is essential for high-fidelity DNA double strand break (DSB) repair and involves numerous genes including BRCA1 and BRCA2. HR deficiency (HRD) due to inactivation of such genes leads to increased levels of genomic alterations[1]. HRD is a common characteristic of many tumors and is frequently observed in breast and ovarian cancer[2]. Accurate detection of HR deficiency (HRD) is of clinical relevance as it is indicative of sensitivity to targeted therapy with poly ADP-ribose polymerase inhibitors (PARPi)[3, 4] as well as to DNA damaging reagents[1].

In the clinic, germline BRCA1/2 mutation status is currently the main genetic biomarker of HRD[5]. However, germline testing has its drawbacks: i) it is dependent on the completeness and accuracy of clinical variant annotation databases (e.g. ClinVar); ii) epigenetic silencing is overlooked; iii) partial/complete deletions of the BRCA1/2 loci are missed by current clinical genetic testing, resulting in BRCA1/2 status reporting based on the wild type allele from contaminating normal tissue; and iv) HRD can be driven purely by somatic events. Furthermore, the focus on BRCA1/2 overlooks inactivation of other HR pathway genes. Consequently, patients may receive incorrect treatment or miss out on treatment opportunities, thus necessitating the development of better biomarkers for HRD.

It was recently shown that somatic passenger mutations, which are identified efficiently by whole genome sequencing (WGS), can provide insights into the mutational processes that occurred before and during tumorigenesis, paving the way for novel opportunities for clinical tumor diagnostics[6]. For the repair of DSBs, HRD tumors are dependent on alternative more error-prone pathways including microhomology mediated end-joining (MMEJ)[7], resulting in a characteristic mutational footprint across the genome that can be used to detect HRD regardless of the underlying cause (whether genetic or epigenetic). Indeed, some mutational footprints were found to be associated with BRCA1/2 deficiency, namely deletions with flanking microhomology, as well as several ‘mutational signatures’ including two COSMIC single nucleotide variant (SNV) signatures and two structural variant (SV) signatures[8]. These features were used to develop a breast cancer-specific predictor of HRD known as HRDetect[9]. Application of this tool in primary tumors revealed that the prevalence of HRD extends beyond BRCA1/2-deficient breast cancer tumors, and occurs at varying frequencies in different cancer types[10]. However, HRD rates in advanced metastatic cancer remain unclear, although these are the patients that are increasingly targeted with personalized treatments including PARP inhibitors for BRCA-deficiency[5].

Here, we describe the development of a random forest-based **C**lassifier of **HO**mologous **R**ecombination **D**eficiency (CHORD) for pan-cancer HRD detection. With this model, we demonstrate that accurate prediction of HRD is possible across cancer types using specific SNV, indel and SV types. We identified inactivation of BRCA1, BRCA2, RAD51C and PALB2 as the most frequent genetic cause of HRD pan-cancer in both primary and metastatic cancer, with the latter two genes resulting in the same mutational footprints as BRCA2 (consistent with the findings of recent studies in breast cancer[11, 12]). In addition, we found that the underlying genetic inactivation of these genes was cancer type specific, but independent of tumor progression state.

## Results

### Random forest classifier training

For the development of CHORD, we used WGS data of 3,824 solid tumors from 3,584 patients from the pan-cancer metastatic cohort of the Hartwig Medical Foundation (HMF)[13]. From these, we selected tumor samples with biallelic loss of BRCA1 or BRCA2, and non-mutated BRCA1/2, to obtain a high confidence set of samples belonging to 3 classes for classifier training (BRCA1-deficient, BRCA2-deficient, and BRCA1/2-proficient). To this end, we screened each sample to identify those samples with one of the following events in BRCA1/2: (i) complete copy number loss (i.e. deep deletion), (ii) loss-of-heterozygosity (LOH) in combination with a pathogenic germline or somatic SNV/indel (as annotated in ClinVar, or a frameshift), or (iii) 2 pathogenic SNV/indels. This unbiased approach revealed 35 and 89 samples with BRCA1 or BRCA2 biallelic loss of function, respectively, which were labeled as HRD for the training. Conversely, 1,902 samples were labeled as HR proficient (HRP) as these samples were observed to carry at least one functional allele of BRCA1/2. In total, 2,026 out of 3,824 samples (53% of the HMF dataset) were used to train the classifier (**Supplementary figure 1**).

The occurrence of three main somatic mutation categories were used as features for training (**Figure 1a**), which included (i) single nucleotide variants (SNVs) subdivided by base substitution (SBS) type; (ii) indels stratified by the presence of sequence homology, tandem repeats, or the absence of either; and (iii) structural variants (SV), stratified by type and length. An initial feature analysis revealed that small deletions with ≥2bp flanking homology were together more predictive of BRCA1/2 deficiency versus deletions with 1bp flanking homology (**Supplementary figure 3**). Thus, deletions with flanking homology were further split into these two homology length bins. The occurrence of the 29 features together formed a contribution profile for each sample. From this, relative contributions per mutation category were calculated to account for differences in mutational load across samples (**Figure 1a**). These features are henceforth collectively referred to as ‘mutation contexts’.

**Figure 1:**
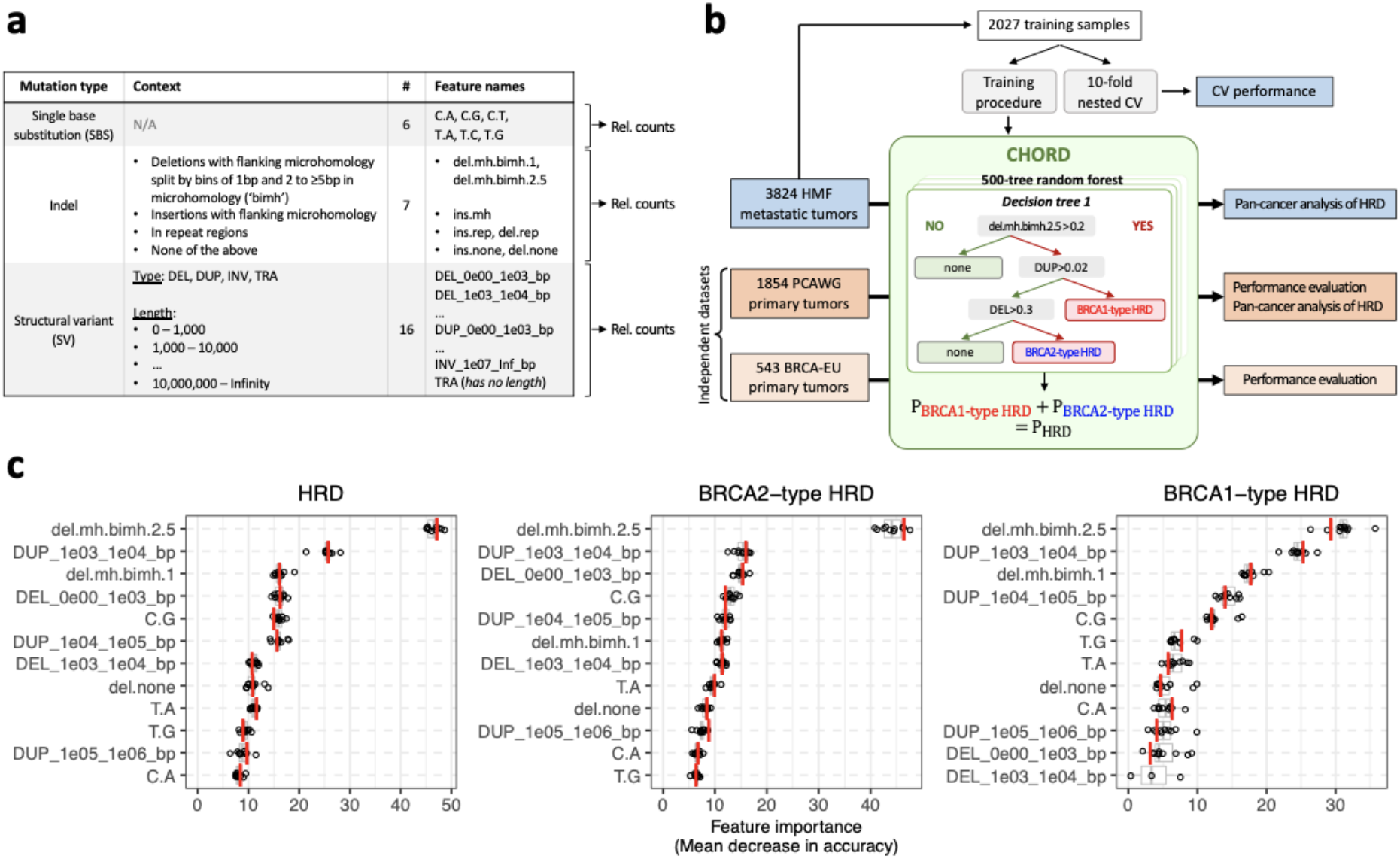
CHORD is a random forest **C**lassifier of **HO**mologous **R**ecombination **D**eficiency able to distinguish between BRCA1- and BRCA2-type HRD phenotypes in a pan-cancer context. (**a**) The features used for training CHORD are relative counts of different mutation contexts which fall into one of three groups based on mutation type. (i) Single nucleotide variants (SNV): 6 possible base substitutions (C>A, C>G, C>T, T>A, T>C, T>G). (ii) Indels: indels with flanking microhomology, within repeat regions, or not falling into either of these 2 categories. (iii) Structural variants (SV): SVs stratified by type and length. Relative counts were calculated separately for each of the 3 mutation types. (**b**) Training and application of CHORD. From a total of 3,824 metastatic tumor samples, 2,026 samples were selected for training CHORD. The model outputs the probability of BRCA1-type HRD and BRCA2-type HRD, with the probability of HRD being the sum of these 2 probabilities. The performance of CHORD was assessed via a 10-fold nested cross-validation (CV) procedure on the training samples, as well as by applying the model to the BRCA-EU dataset (543 primary breast tumors) and PCAWG dataset (1,854 primary tumors). Lastly, CHORD was applied to all samples in the HMF and PCAWG dataset in order to gain insights into the pan-cancer landscape of HRD. (**c**) The features used by CHORD to predict HRD as well as BRCA1-type HRD and BRCA2-type HRD, with their importance indicated by mean decrease in accuracy. Deletions with 2 to ≥5bp (i.e. ≥2bp) of flanking microhomology (del.mh.bimh.2.5) was the most important feature for predicting HRD as a whole, with 1-100kb structural duplications (DUP_1e03_1e04_bp, DUP_1e04_1e05_bp) differentiating BRCA1-type HRD from BRCA2-type HRD. Boxplot and dots show the feature importance over 10-folds of nested CV on the training set, with the red line showing the feature importance in the final CHORD model. Boxes show the interquartile range (IQR) and whiskers show the largest/smallest values within 1.5 times the IQR.

A random forest was then trained to predict the probability of BRCA1 or BRCA2 deficiency (**Figure 1b**). Briefly, a core training procedure performed feature selection and class resampling (to alleviate the imbalance between the 3 classes). This core procedure was subjected to 10-fold cross-validation (CV) which was repeated 100 times to filter samples from the training set that were not consistently HRD or HRP. A sample was considered HRD if the sum of the BRCA1 and BRCA2 deficiency probabilities (henceforth referred to as the HRD probability) was ≥0.5. This core procedure was reapplied to the filtered training set to yield the final random forest model which we refer to as ‘CHORD’ (**Supplementary figure 2a,b**; **Supplementary figure 4**).

The presence of deletions with ≥2bp flanking homology (del.mh.bimh.2.5) was found to be the most important predictor of HRD. Additionally, CHORD uses 1-10kb and to a lesser extent 10-100kb duplications (DUP_1e03_1e04_bp and DUP_1e04_1e05_bp, respectively) for distinguishing BRCA1 from BRCA2 deficiency. Given that deficiencies in other HR genes may lead to similar phenotypes, we have coined the terms ‘BRCA1-type HRD’ and ‘BRCA2-type HRD’ to describe these HRD subtypes (**Figure 1c**). Together, the features that are predictive of HRD are in line with those of a previously described HRD classifier HRDetect^9^. However, the feature weights differ markedly likely due to differences in the background mutational landscape between the pan-cancer cohort used here compared to the breast cancer cohort used for training HRDetect.

### Performance of CHORD

Two approaches were used to assess the performance of CHORD. In the first approach, 10-fold CV was performed on the training data which allows every sample to be excluded from the training set after which unbiased HRD probabilities can be determined (**Supplementary figure 2c**). The probabilities of all prediction classes (i.e. HRD, BRCA1-type HRD, BRCA2-type HRD) were highly concordant with the genetic annotations (**Figure 2a**). The concordance between predictions and annotations was quantified by calculating the area under the curve of receiver operating characteristic (AUROC) and precision-recall (AUPRC) curves (**Figure 2b,c**). CHORD achieved excellent performance as shown by the high AUROC and AUPRC for all prediction classes (0.98 and 0.87 respectively). Additionally, CHORD achieved a maximum F1-score (~0.88) for predicting HRD at a cutoff of 0.5 which was thus set to be the classification threshold (**Supplementary figure 6**).

**Figure 2:**
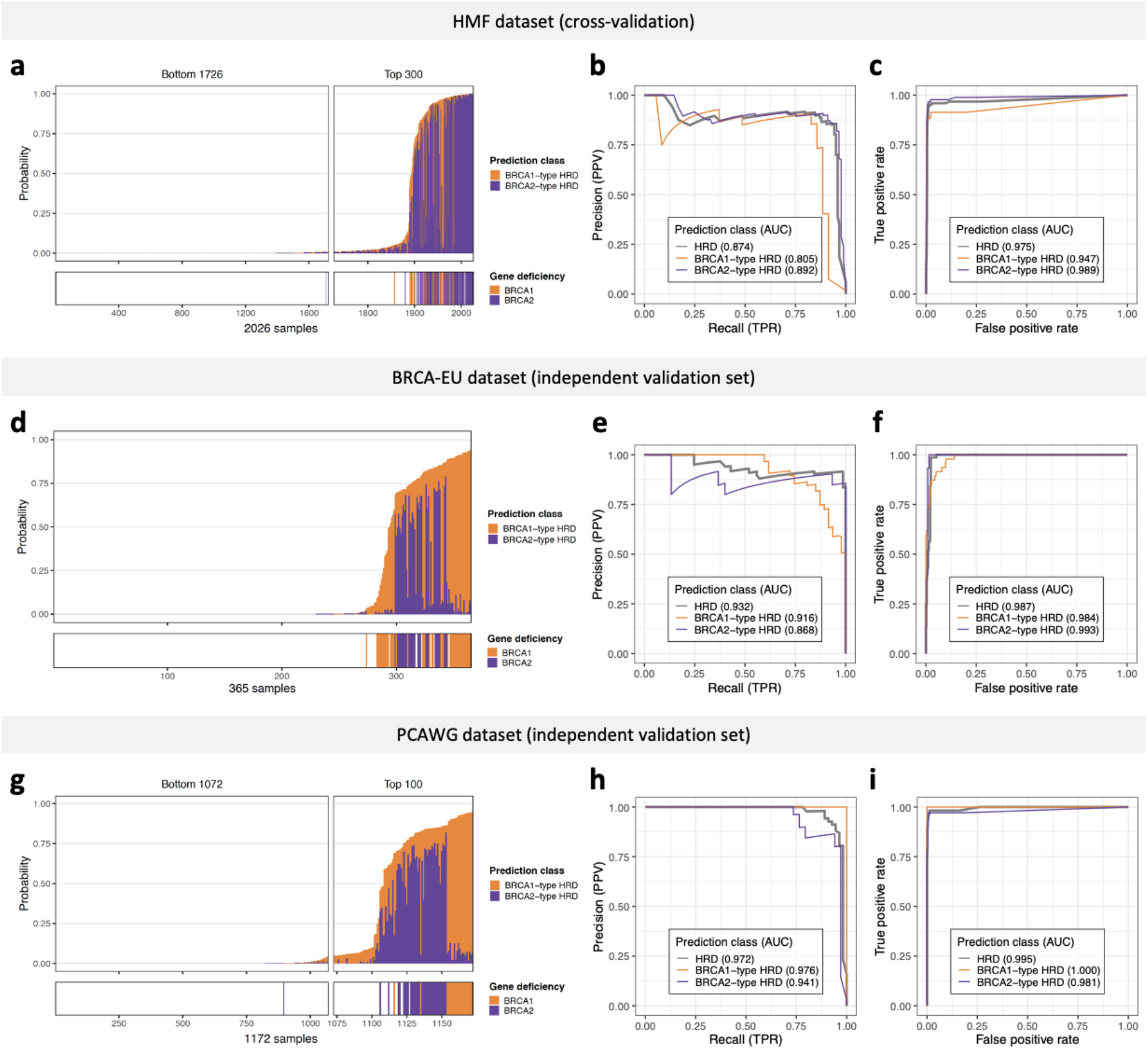
Performance of CHORD as determined by 10-fold cross-validation (CV) on the HMF training data or prediction on two independent datasets: BRCA-EU (primary breast cancer dataset) and PCAWG (primary pan-cancer dataset). BRCA-EU and PCAWG samples shown here all passed CHORD’s QC criteria (i.e. MSI absent, ≥50 indels, ≥30 SVs if a sample was predicted HRD). (**a, d, g**) The probability of HRD for each sample (total bar height) with each bar being divided into segments indicating the probability of BRCA1- (orange) and BRCA2-type HRD (purple). Stripes below the bar plot indicate biallelic loss of BRCA1 or BRCA2. In (**a**), probabilities have been aggregated from the 10 CV folds. (**b, e, h**) Receiver operating characteristic (ROC) and (**c, f, i**) precision-recall curves (PR) and respective area under the curve (AUC) values showing the performance of CHORD when predicting HRD as a whole (grey), BRCA1-type HRD (orange), or BRCA2-type HRD (purple).

In the second approach, performance was evaluated on two independent datasets: the BRCA-EU dataset[8] (543 primary breast tumors) and the PCAWG dataset[14] (1,854 primary tumors, pan-cancer). For both datasets, samples that (i) passed CHORD’s QC criteria (i.e. MSI absent, ≥50 indels, ≥30 SVs if a sample was predicted HRD; **Supplementary information**) and (ii) for which the biallelic status of BRCA1/2 could confidently be determined were selected for validation of CHORD. For the BRCA-EU dataset, this included the 365 samples that were used to train and evaluate the performance of HRDetect[9]. For the PCAWG dataset, this included 1,172 samples for which the same genetic criteria used for selecting samples from the HMF dataset for training CHORD applied. Applying CHORD on these samples revealed that the HRD probabilities were concordant to their BRCA1/2 genetic status for both the BRCA-EU and PCAWG datasets (**Figure 2d,g**). The AUROC (>0.98) and AUPRC (>0.93) values were comparable to those obtained by CV on the HMF training data for all prediction classes for both datasets (**Figure 2e,f,h,i**). In the BRCA-EU dataset, we still observed some BRCA1 deficient samples classified as HRP by CHORD (while HRDetect classified these as HRD) and tested whether this was due to differences in somatic calling algorithms. Indeed, using the variants obtained from the native pipeline of the HMF dataset (HMF pipeline[13]) for HRD prediction resulted in overall higher HRD probabilities compared to using the variants downloaded from ICGC, especially for BRCA1-deficient samples. This was apparent for sample PD4017 which became HRD using HMF pipeline called mutation profiles, with PD24186, PD11750 and PD23578 having greatly increased HRD probabilities (**Supplementary figure 7**). Our results thus demonstrate that CHORD is robust when applied to other datasets. However, differences in variant calling pipelines can affect CHORD’s ability to predict HRD (especially considering the still existing challenges of indel and SV calling from WGS data, and CHORD’s dependency on these features). Additional validation and threshold optimisation is thus recommended when applying CHORD on data from other variant calling pipelines.

We note that CHORD performs similarly to HRDetect based on predictions on the BRCA-EU dataset (AUROC=0.98 for both models)[9]. Additionally, the predictions of CHORD and HRDetect on the PCAWG dataset[10] were concordant for the vast majority of samples (1506/1526; 99%) (**Supplementary figure 8**). Of the 8 HRD samples only detected by CHORD, 3 showed biallelic loss of BRCA1/2, while none of the HRDetect-only samples could be explained by genetic biallelic loss. Given that CHORD, unlike HRDetect, does not rely on refitting algorithms on SBS signatures[6] and SV signatures[8], the similar performance between the two models suggests that accurate detection of HRD is possible without using mutational signatures. To further validate this, we also trained a random forest model (CHORD-signature) that uses the SBS and SV signatures as input instead of mutation contexts. CHORD-signature performed similarly to CHORD (**Supplementary figure 10**), which can be explained by the reliance on similar features (**Supplementary figure 9**), namely microhomology deletions and SV signature 3 (analogous to 1-100kb duplications). SBS signature 3 (proposed as a sensitive marker for HRD in recent studies[15–17]) is actually a less important feature for predicting HRD than microhomology deletions in both HRDetect (as demonstrated by the authors[9]) and CHORD-signature, indicating that microhomology deletions serves as a better (univariate) marker of HRD compared to SBS signature 3 [10]. We thus conclude that accurate detection of HRD does not require mutational signatures, thereby simplifying HRD calling and avoiding the complications associated with the fitting step required for computing signature contributions in individual samples (for which there is currently no consensus approach)[18].

### Effect of treatment on HRD predictions

The HMF dataset comprises tumors from patients with metastatic cancer who have been exposed (some heavily) to treatment which could potentially affect CHORD’s predictions. Two recent studies showed that common cancer treatments in general do not induce mutations that may interfere with CHORD predictions[19, 20]. However, these two studies (as well as one by Behjati *et al.* [21]) did show that radiotherapy had the potential to induce deletions with flanking microhomology, which could potentially lead to false positive HRD classifications. To investigate this, we used random forests to identify and compare the mutational features associated with radiotherapy and BRCA1/2 deficiency when using clonal variants versus subclonal variants (which are enriched for treatment induced mutations[19, 22]) as input features. This revealed that small deletions with 1bp of flanking homology (del.mh.bimh.1) are highly associated with radiotherapy (**Supplementary figure 12**) and less with BRCA1/2 deficiency. Also, when we retrain CHORD with all microhomology deletions merged into a single feature (CHORD-del.mh.merged; **Supplementary figure 13**), we observed a higher number of samples as being HRD based on subclonal variants but HRP based on clonal variants (97) compared to CHORD (64 samples) (**Supplementary figure 14a,b**). This indicates that having the microhomology deletions feature split by these two homology length bins may mitigate radiotherapy-associated false positive predictions. Importantly, some samples that are scored HRP based on clonal variants but HRD on subclonal variants could truly be HRD, especially since 4 of these samples had evidence of BRCA1/2 biallelic loss (deep deletion: n=1; LOH and a pathogenic variant: n=1; 2 pathogenic variants: n=2). For these samples, it is likely that BRCA1/2 biallelic loss occurred relatively late in the tumor progression stage which results in an insufficient number of HRD-associated mutations for clear HRD classification by CHORD. Furthermore, subclonal-only HRD could potentially also be explained by transient inactivation of HR e.g. through epigenetic silencing of key components. Nevertheless, as treatment induced effects can formally never be excluded, CHORD predictions on subclonal variants must in any case be interpreted with caution, especially given the extra challenges associated with accurately detecting subclonal variants with low variant allele frequency (VAF).

Overall, a similar number of samples were predicted HRD for both CHORD and CHORD-del.mh.merged when using clonal variants (128 and 127 out of 1,724; **Supplementary figure 14a,b**), and these numbers were also similar to when using all variants (135 and 132; **Supplementary figure 14c,d**), indicating that radiotherapy likely has minimal impact on HRD predictions when not using subclonal variants as input.

### BRCA2, RAD51C and PALB2 are associated with BRCA2-type HRD while only BRCA1 is associated with BRCA1-type HRD

To gain insights into the genetic causes of HRD, we applied CHORD to both the HMF and PCAWG datasets and selected the samples that passed CHORD’s QC criteria (i.e. MSI absent, ≥50 indels, ≥30 SVs if a sample was predicted HRD; **Supplementary information**). For the HMF dataset, we also selected a single tumor per patient (based on highest tumor purity) for those with multiple biopsies, though all patients had consistent HRD probabilities across all biopsies (**Supplementary table 1**). This yielded a total of 5,122 patients (3,504 from HMF and 1,618 from PCAWG), with 310 (6%) being classified as being homologous recombination deficient (CHORD-HRD). Of these, 121 were classified as having BRCA1-type HRD and 189 as having BRCA1-type HRD. The remaining 4,812 patients were classified as homologous recombination proficient (CHORD-HRP) (**Figure 3a**, **Supplementary table 1**).

**Figure 3:**
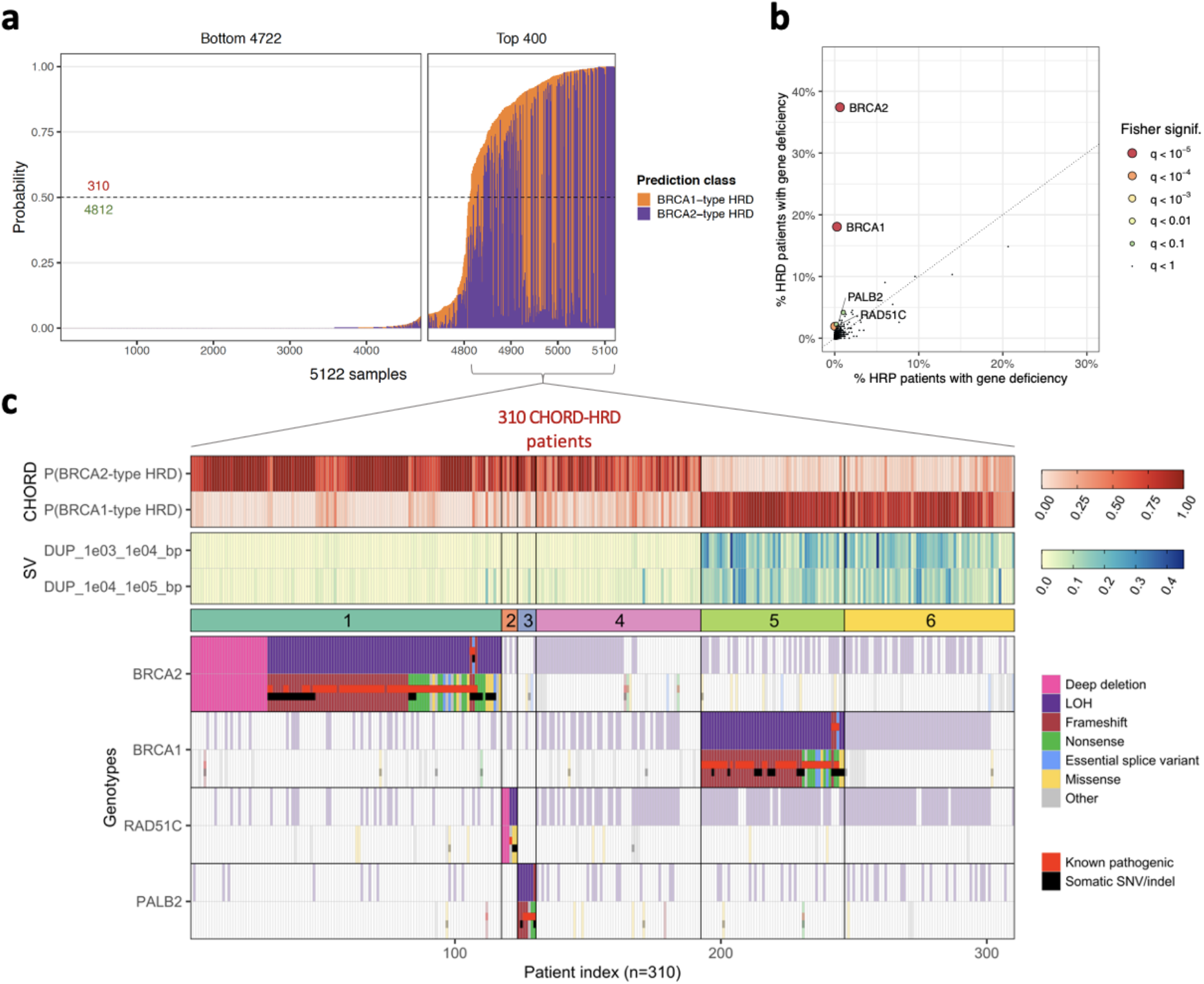
The genetic causes of HRD in patients from the HMF and PCAWG datasets. (**a**) The bar plot shows the probability of HRD for each patient (total bar height) with each bar being divided into segments indicating the probability of BRCA1-type HRD (orange) and BRCA2-type HRD (purple). 310 patients were predicted HRD while 4,812 were predicted HRP by CHORD. (**b**) A one-tailed Fisher’s exact test identified enrichment of BRCA1, BRCA2, RAD51C and PALB2 biallelic inactivation in CHORD-HRD vs. CHORD-HRP patients (from a list of 781 cancer and HR related genes). Each point represents a gene with its size/color corresponding to the statistical significance as determined by the Fisher’s exact test, with axes indicating the percentage of patients (within either the CHORD-HRD or CHORD-HRP group) in which biallelic inactivation was detected. Multiple testing correction was performed using the Hochberg procedure. (**c**) Biallelic inactivation of BRCA2, RAD51C and PALB2 was associated with BRCA2-type HRD, whereas only BRCA1 inactivation was associated with BRCA1-type HRD. Top: BRCA1- and BRCA2-type HRD probabilities from CHORD. Middle: SV contexts used by CHORD to distinguish BRCA1-from BRCA2-type HRD. Bottom: The biallelic status of each gene. Samples were clustered according to HRD subtype, and by the impact of a biallelic/monoallelic event (based on ‘P-scores’ as detailed in the methods). Clusters 1, 2, 3, and 5 correspond to patients with identified inactivation of BRCA2, RAD51C, PALB2 and BRCA1, while clusters 4 and 6 correspond to patients without clear biallelic inactivation of these 4 genes. Tiles marked as ‘Known pathogenic’ refer to variants having a ‘pathogenic’ or ‘likely pathogenic’ annotation in ClinVar. ‘Other’ variants include various low impact variants such as splice region variants or intron variants (these are fully specified in **Supplementary table 4**). Only data from samples that passed CHORD’s QC criteria are shown in this figure (MSI absent, ≥50 indels, and ≥30 SVs if a sample was predicted HRD).

We then sought to identify the key mutated genes underlying the HRD phenotype by performing an enrichment analysis of biallelically inactivated genes in CHORD-HRD vs. CHORD-HRP patients. For this analysis, we started from a list of 781 genes that are cancer related (based on the catalog of genes from Cancer Genome Interpreter) and/or HR related (manually curated based on the KEGG HR pathway, as well as via literature search) (**Supplementary table 3**). For these genes, we considered likely pathogenic variants (according to ClinVar) as well as predicted impactful variants such as nonsense mutations to contribute to gene inactivation (see *Methods*). This revealed that, in addition to BRCA1 and BRCA2 (q<10^−5^ for both genes, one sided Fisher’s exact test), RAD51C and PALB2 (q<0.001 and q<0.05 respectively) were also significantly enriched amongst HRD patients using a q-value threshold of 0.05 (**Figure 3b**).

Of all CHORD-HRD HMF patients, ~60% (184/310) could be explained by biallelic inactivation of either BRCA2 (cluster 1; n=117), BRCA1 (cluster 5; n=54), RAD51C (cluster 2; n=6), or PALB2 (cluster 3; n=7), which was most often caused by LOH in combination with a pathogenic variant or frameshift, or a deep deletion (**Figure 3c**, **Supplementary table 4**). RAD51C and PALB2 were recently linked to HRD as incidental cases using mutational signature based approaches[11, 16] and our results now confirm that biallelic inactivation of these two genes results in HRD and is actually a common cause of HRD (albeit to a lesser extent than for BRCA1/2). RAD51C and PALB2 deficient patients shared the BRCA2-type HRD phenotype (absence of duplications) with BRCA2 deficient patients (clusters 1-3; **Figure 3c**), consistent with previous studies[11, 12]. On the other hand, only BRCA1 deficient patients (cluster 5) harbored the BRCA1-type HRD phenotype (1-100kb duplications).

Of note, we observed one patient (**Figure 3c**; patient #6) bearing a known pathogenic frameshift mutation in BRCA1 (**Supplementary table 4**; patient HMF001925, c.1961dupA), which based on current practices for detecting HRD in the clinic (testing for pathogenic SNVs/indels)[5] would be considered the driver mutation. However, our genetic analysis indicates that the deep deletion in BRCA2 (which would be missed by testing for SNVs/indels) was the cause of HRD, which is supported by the lack of LOH in BRCA1, as well as the BRCA2-type HRD phenotype of this patient.

In ~40% of CHORD-HRD patients (126/310; clusters 4 and 6, **Figure 3c**), there was no clear indication of biallelic loss of BRCA1/2, RAD51C or PALB2 (henceforth referred to as the ‘HRD associated genes’). However, 109 of these patients had a deleterious event in a single allele of one of the HRD associated genes (the majority due to LOH (including copy number neutral LOH)), with a similar cancer type distribution in these patients as in the biallelically affected patients (**Supplementary figure 18**). We found enrichment of LOH in BRCA1, BRCA2, as well as RAD51C in HRD samples (**Supplementary figure 19**), which implies the involvement of LOH in the inactivation of these genes for the patients in clusters 4 and 6. This is consistent with the finding by Jonsson et al.[15] that LOH is enriched in tumors with BRCA1/2 germline pathogenic variants or somatic loss-of-function variants. Davies et al.[9] showed that promoter methylation of BRCA1 was present in 22% of ovarian and 16% of breast primary cancers with HRD (**Table 1**). BRCA1 and RAD51C promoter methylation with loss of the other allele was also reported in HRD tumors in other studies[11, 12, 16]. Thus, BRCA1 and RAD51C promoter methylation, likely in combination with LOH, may have led to the HRD phenotype for a sizable portion of the ovarian and of breast cancer patients with no clear biallelic loss of the HRD associated genes, and potentially for patients with other cancer types as well (**Supplementary figure 20**). Unfortunately, we could not directly assess this as methylation data was not available for the HMF nor the PCAWG dataset.

**Table 1:** **(A)** The number of ovarian and breast cancer HRD patients with BRCA1 promoter methylation as reported by Davies et al.[9] (from supplementary tables 1, 3, and 8) in comparison to **(B)** the number of CHORD-HRD ovarian and breast cancer patients without biallelic loss of the HRD associated genes (BRCA1/2, RAD51C, PALB2).

| Cancer type | (A) HRD patients with BRCA1 promoter methylation (Davies et al.) |  | (B) CHORD-HRD patients without biallelic loss of HRD associated genes |  |
| --- | --- | --- | --- | --- |
|  | % | # / total | % | # / total |
| Ovarian | 22% | 10/46 | 50% | 21/42 |
| Breast | 16% | 14/86 | 49% | 38/77 |

We also cannot rule out the possibility that deficiencies in other HR genes that did not reach significance in our enrichment analysis, underlie the HRD phenotype for a small number of patients in clusters 4 and 6. We indeed identified 17 patients with biallelic inactivation of a HR gene other than BRCA1/2, RAD51C or PALB2, and 1 patient with a likely inactivating biallelic event (LOH in combination with a nonsense variant in CHEK1) (**Supplementary figure 21**). Notably, the 4 patients with RAD51B (n=2) and XRCC2 (n=2) deficiency were all predicted to have BRCA2-type HRD, a phenotype shared with RAD51C deficient patients[23]. Given that these 3 genes all belong to the RAD51 paralog complex BCDX2[24], the BRCA2-type HRD suggests that RAD51B and XRCC2 deficiency could have led to HRD in these patients. Likewise, the 4 patients with deficiencies in the BRCA1-binding proteins, BARD1[25] (n=1), BRIP1[26] (n=1), FAM175A[27] (n=1) and FANCA[28] (n=1), were all predicted as having BRCA1-type HRD. Thus, while we could not conclusively determine the cause of HRD for patients in clusters 4 and 6, we postulate that HRD in these patients may have been a result of epigenetic silencing of BRCA1/2 or RAD51C, deficiencies in other HR genes (not associated to HRD in our analysis), or possibly a result of other unknown regulatory mechanisms.

### The incidence and genetic cause of HRD varies in different tissue types and cancer stage

We next investigated the differences in the incidence and genetic causes of HRD based on primary tumor location in both primary (PCAWG) and metastatic (HMF) cancer datasets (**Figure 4**). HRD was most prevalent in ovarian, breast, prostate and pancreatic cancer (85% combined), and only occurred sporadically in other cancer types (15%) (**Supplementary table 5**). Compared to metastatic cancer, HRD is found much more often in primary ovarian (52% vs 30%) and breast (24% vs 12%) cancers, and less often in primary prostate (5.6% vs 13%) and pancreatic (7.3% vs 13%) cancer (**Figure 4a**). Notably, in metastatic cancer, prostate and pancreatic cancer has a similar incidence of HRD to breast cancer (all ~13%). However, the observed differences in HRD rates between the primary and metastatic cohorts may not necessarily be conclusive as we can not rule out confounding factors such as patient inclusion criteria.

**Figure 4:**
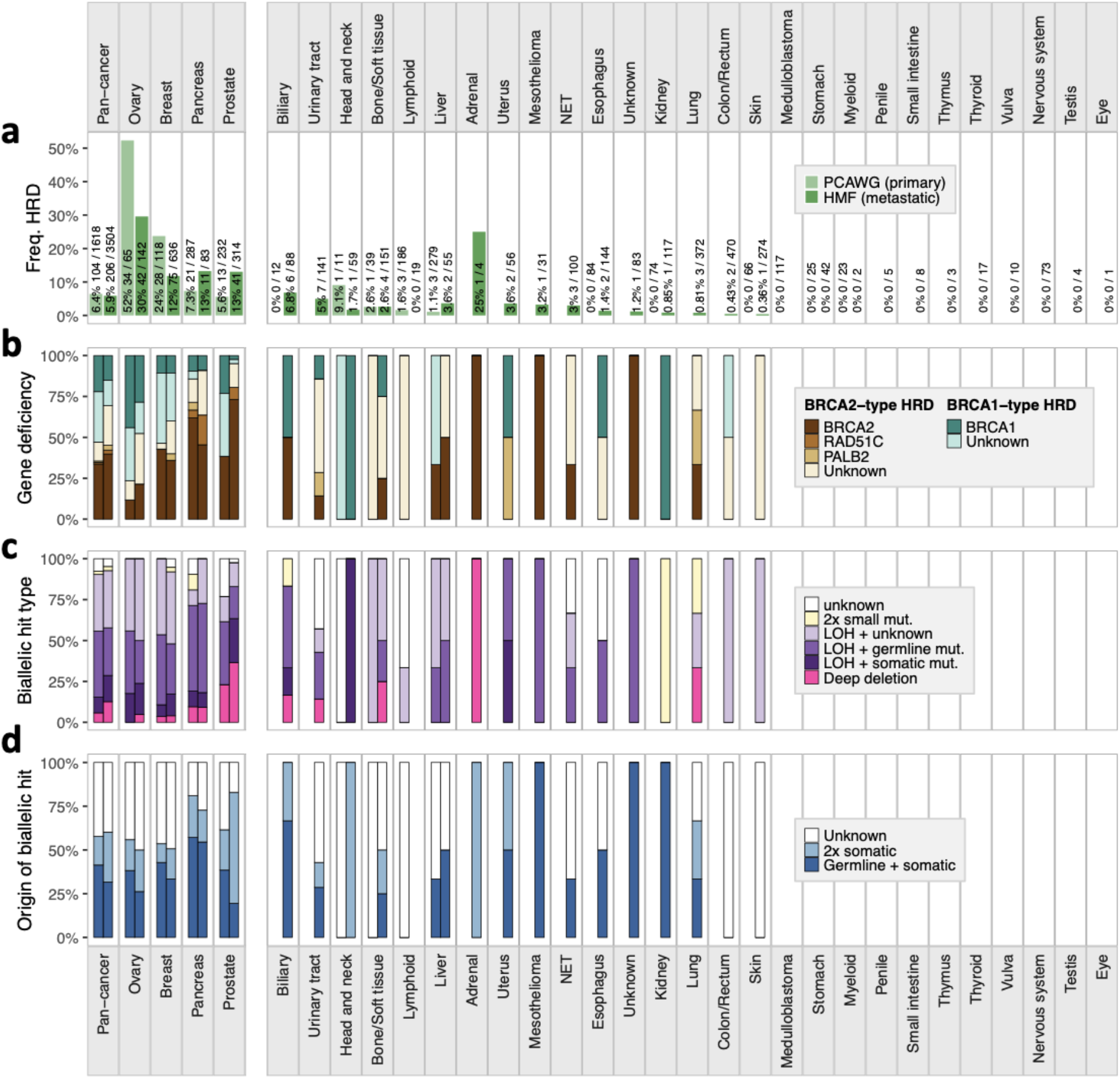
Percentage breakdown of the incidence and genetic causes of HRD in CHORD-HRD patients pan-cancer and by cancer type in the HMF and PCAWG datasets (left and right bars respectively). The vertical split in the figure separates cancer types with (left side) and without (right side) ≥10 CHORD-HRD patients in at least one of the datasets. (**a**) Frequency of HRD. Cancer types where no frequency of HRD is displayed contain no data in either the HMF or PCAWG datasets (**b**) The gene deficiency associated with HRD. Bar segments are grouped into BRCA2-type HRD genes (BRCA2, RAD51C, PALB2) and BRCA1-type HRD genes (BRCA1 only). (**c**) The likely combination of biallelic events in BRCA1/2, RAD51C or PALB2 causing HRD. (**d**) Whether the genetic cause of HRD was purely due to somatic events, due to germline predisposition, or unknown. In (**c** and **d**), ‘Unknown’ and/or ‘LOH + unknown’ bar segments refer to patients where no clear biallelic loss of the aforementioned BRCA1/2, RAD51C, or PALB2 was identified (i.e. clusters 4 and 6 of **Figure 3c**). Only data from samples that passed CHORD’s QC criteria are shown in this figure (MSI absent, ≥50 indels, and ≥30 SVs if a sample was predicted HRD).

Across different cancer types, we observed pronounced diversity in HR function loss (**Figure 4b**). BRCA2-type HRD deficiencies (including BRCA2, RAD51C, PALB2 deficiencies) were more frequent in pancreatic and prostate cancer. On the other hand, BRCA1-type HRD deficiencies were found more often in ovarian and breast cancer. Interestingly, for ovarian and prostate cancer, BRCA1-type HRD deficiencies were more prominent in primary cancer compared to metastatic cancer. Whether these differences in gene deficiencies in different cancer types can be linked to a biological cause or have prognostic value remains to be determined.

In 94% (292/310) of all CHORD-HRD patients, we found mono- or biallelic inactivation of at least one of the four HRD associated genes (BRCA1, BRCA2, PALB2, RAD51C; **Figure 4c**). In the case of biallelic inactivation, we observed LOH to be the dominant secondary event, occurring in combination with a germline SNV/indel (33%) or with a somatic SNV/indel (14%) of CHORD-HRD patients. LOH of BRCA1/2, RAD51C or PALB2 was also found as a monoallelic event, mainly in ovarian (47%) and breast (49%) cancer patients (**Supplementary table 5**). As indicated earlier, the other allele may be inactivated by epigenetic mechanisms in these patients (or alternatively HRD was caused by inactivation of another HR gene). Interestingly, we find that deep somatic deletions do frequently contribute to biallelic loss of BRCA2 or RAD51C, occurring in 10% of CHORD-HRD patients pan-cancer (**Supplementary table 5**). However, deep deletions (primarily of BRCA2; **Supplementary figure 20**) occurred much more frequently in prostate cancer (33%) compared to other cancer types, consistent with previous observations[29]. Nevertheless, deep deletions of HRD genes did occur in every cancer type with a high frequency of CHORD-HRD patients indicating that complete somatic gene loss is a common and underestimated cause of HRD in both primary and metastatic cancer.

We find that biallelic gene loss is often associated with germline predisposition (**Figure 4d**) in ovarian (32%), breast (36%), and pancreatic (56%) cancer patients, but to a lesser extent in prostate cancer patients (24%) (**Supplementary table 5**). On the other hand, biallelic gene loss exclusively by somatic events occurs in sizable proportion of CHORD-HRD patients (35% pan-cancer), being most frequent in prostate cancer (54%) (**Supplementary table 5**) mainly due to the deep deletions (**Supplementary figure 20**). Although these frequencies may not be fully representative for each cancer type due to the proportion of patients with unknown mutation status in at least one allele (indicated as ‘Unknown’ in **Figure 4d**), these observations do emphasize that somatic-only events should not be overlooked as a mechanism of HR gene inactivation.

## Discussion

Here we describe a classifier (CHORD) that can detect HRD (as well as HRD sub-phenotypes) across cancer types based on mutation profiles. By using this tool in a systematic pan-cancer analysis, we reveal novel insights into the mechanisms and incidence of HRD across cancer types with potentially important clinical relevance.

HRD targeted therapy with PARPi is mostly restricted to breast and ovarian cancer[5], though its use for treating pancreatic cancer was recently approved by the FDA (US Food and Drug Administration)[30]. However, we show that HRD is common not only in ovarian and breast cancer, but also in prostate, pancreatic cancer. The incidence of HRD was relatively higher in metastatic prostate and pancreatic cancer, and lower for ovarian and breast cancer as compared with primary tumors. This may reflect more intensive familial (germline) testing for BRCA1/2 mutations in ovarian and breast cancer[31] and consequently earlier diagnosis and treatment with fewer cases of progression to metastatic cancer as a result. However, we cannot formally exclude that these observations originate from differences in cohort inclusion criteria that could skew numbers (e.g. due to more recruitment of patients with triple negative breast cancer which has higher HRD rates[12]).

We show that HRD is also found sporadically in cancer types beyond breast, ovarian, prostate or pancreatic, but collectively this constitutes a sizable group of patients (15% of all patients). Our results thus indicate that a large number of patients who would potentially benefit from PARPi therapy still remain unnoticed. Since the mutational phenotype of HRD is independent of cancer type, mutational scar based HRD detection such as with CHORD would be valuable for cancer type agnostic patient stratification for future PARPi trials[32]. This is particularly important for metastatic patients (who depend on systemic treatments and benefit most from targeted treatments like PARPi), as well as for cancer types currently lacking good markers for patient stratification for such treatment (such as prostate[33] and biliary[34] cancer).

Genetic based detection of HRD in the clinic is commonly done by testing for pathogenic BRCA1/2 germline mutations[5]. However, such hereditary mutations are only present in 30% of CHORD-HRD patients (**Supplementary information**) indicating that germline testing likely misses a substantial number of HRD patients. Germline variant testing is particularly unsuitable for prostate cancer where gene inactivation is frequently caused by somatic deep deletions, which prevent the identification of any SNVs/indels at the affected locus when using panel- or PCR-based sequencing methods (exon scanning). This problem also exists for other cancer types where deep deletions also make up a non-negligible fraction of HR gene inactivation cases. While somatic mutation testing improves diagnostic yield and is indeed increasingly performed in the clinic[5], WGS based genetic testing is ultimately necessary to capture the full spectrum of genetic alterations and to accurately determine the mutational status of HR genes. However, even such broad genetic testing with focus on biallelic gene inactivation still potentially misses roughly 50% of all HRD patients (**Supplementary information**).

We do acknowledge that mutational scars represent genomic history and not current on-going mutational processes that can result in false positive CHORD predictions, which could be for example due to reversion of HRD by secondary frameshifts[35, 36], or recent acquisition of the HRD phenotype. False positive predictions could also arise from treatments producing similar mutational scars (in particular, microhomology deletions) to HRD. The most common cancer treatments have been shown to have little or no contribution to microhomology deletions, with the exception of radiotherapy[19–21]. However, we showed that radiotherapy itself likely does not lead to false positive predictions. We cannot exclude the possibility however that clonal expansion of a radiotherapy resistant tumor cell leads to sufficient enrichment of radiotherapy associated microhomology deletions in the tumor, resulting in a false positive prediction. Ultimately, the ability for CHORD to improve patient stratification and treatment outcome will need to be evaluated in direct comparisons and prospective clinical trials.

Thus, while CHORD can detect HRD independent of the underlying cause, genetic testing of HRD genes is complementary and can provide supporting information for making a final verdict on a patient’s HR status. The unique advantage of using WGS, although not routine in clinical diagnostics yet, but likely in the near future[37], is that both genetic testing and mutational scar based HRD detection with CHORD can be performed simultaneously with the same assay. We envision that the findings from our analyses incentivizes improvements to current clinical practices for detecting HRD, and that the application of genomics-based approaches, like CHORD, in the clinic will support these endeavors and provide additional treatment options for patients. CHORD is freely available as an R package at https://github.com/UMCUGenetics/CHORD.

## Acknowledgements

This publication and the underlying study have been made possible partly on the basis of the data that Hartwig Medical Foundation and the Center of Personalised Cancer Treatment (CPCT, The Netherlands) have made available to the study. We thank Neeltje Steeghs (Netherlands Cancer Institute), Martijn Lolkema (Erasmus Medical Center Rotterdam), Geert Cirkel (Meander Medical Center), Els Witteveen (UMC Utrecht), Mariette Labots (Amsterdam UMC, location VUmc) and Laurens Beerepoot (Elisabeth-TweeSteden Ziekenhuis, Tilburg) for study inclusion of a significant part of the patients that were used in this study and Peter Bouwman (Netherlands Cancer Institute, Amsterdam) for critically reading the manuscript. This work was financially supported by the gravitation program CancerGenomiCs.nl from the Netherlands Organisation for Scientific Research (NWO) and Oncode Institute to E.C.

## Methods

### Datasets

We have used patient data of the CPCT-02 (NCT01855477) and DRUP (NCT02925234) clinical studies that are sequenced and uniformly analyzed by the Hartwig Medical Foundation (HMF; https://www.hartwigmedicalfoundation.nl/en/appyling-for-data/). The data transfer agreement (Data Request 10 and 47) were approved by the medical ethical committees (METC) of the University Medical Center Utrecht. We received germline and somatic VCF files of the 3,824 metastatic tumor samples from 3,584 patients in May 2019. For patients with multiple biopsies that were taken at different timepoints, patient IDs were suffixed by ‘A’ for the first biopsy, ‘B’ for the second biopsy, etc (e.g. HMF001423A, HMF001423B). A detailed description of the and the whole patient cohort has been described in detail in Priestley *et al.* 2019[13].

Somatic variant TSV files of the 560 breast cancer (BRCA-EU) dataset were downloaded from the International Cancer Genome Consortium (ICGC; https://dcc.icgc.org/) in August 2017, with BRCA1/2 status annotations for this dataset being obtained from the supplementary data in Davies et al.^9^.

Somatic variant VCF files and somatic copy-number TSV files for the Pan-Cancer Analysis of Whole Genomes (PCAWG) dataset were downloaded from https://dcc.icgc.org/releases/PCAWG on March 3, 2020. PCAWG access for germline data has been granted via the Data Access Compliance Office (DACO) Application Number DACO-1050905 on October 6, 2017 and via https://console.cancercollaboratory.org download portal on December 4, 2017. Germline VCF files were downloaded from the cancer collaboratory download portal on March 21, 2018.

### Variant calling

Variant calling in the HMF dataset was performed previously by HMF (https://github.com/hartwigmedical/pipeline)[13]. Briefly, reads were mapped to GRCh37 using BWA-MEM v0.7.5a with duplicates being marked for filtering. Indels were realigned using GATK v3.4.46 IndelRealigner. GATK Haplotype Caller v3.4.46 was used for calling germline variants in the reference sample. For somatic SNV and indel variant calling, GATK BQSR3 was first used to recalibrate base qualities, followed by Strelka v1.0.14 for the variant calling itself. Somatic SV calling was performed using GRIDSS v1.8.0. Copy-number calling was performed using PURity & PLoidy Estimator (PURPLE), that combines B-allele frequency (BAF), read depth and structural variants to estimate the purity and copy number profile of a tumor sample[38] as well as VAF and clonality (either clonal, subclonal or inaccurate) estimates of each variant.

### Determining gene biallelic status

For samples in the HMF and PCAWG cohorts, biallelic status was determined for 781 genes (**Supplementary table 3**) which included genes associated with cancer, according to Cancer Genome Interpreter (https://www.cancergenomeinterpreter.org/genes), as well as a manually curated set of genes involved in HR (based on the KEGG HR pathway (https://www.genome.jp/), as well as via a literature search). This was performed using an in-house pipeline that interprets copy-number, and germline and somatic SNV/indel data from the HMF variant calling pipeline to determine biallelic gene status (https://github.com/UMCUGenetics/hmfGeneAnnotation).

First, the copy number status in the gene region was determined. If the minimum copy number was <0.3, the gene was considered to have a deep deletion (and by default biallelically inactivated). Else, the gene was screened for 2 mutation events, which included following combinations: (i) loss-of-heterozygosity (LOH) with a germline or somatic SNV/indel; (ii) a germline and somatic SNV/indel; or (iii) 2 somatic SNV/indels.

LOH was considered pathogenic and was automatically given a P-score of 5. LOH occurred if the minimum minor allele copy number within a gene region was <0.2.

Pathogenicity of SNVs/indels was assessed based on pathogenicity annotations from ClinVar (https://www.ncbi.nlm.nih.gov/clinvar/; GRCh37, database date 2020-02-24). For variants without an entry in ClinVar, pathogenicity was assessed based on variant type as determined by SnpEff (http://snpeff.sourceforge.net/; v4.1h). Briefly, variants can be given one of the following annotations from ClinVar: pathogenic, likely pathogenic, variant of unknown significance (VUS), likely benign, and benign. A pathogenicity score (P-score) of 1-5 was also assigned to each annotation, with 1=benign and 5=pathogenic. Additionally, variant types as determined by SnpEff were assigned similar annotations and scores: out-of-frame frameshifts were considered pathogenic (P-score=5); nonsense and splice variants were considered likely pathogenic (P-score=4); missense variants, essential splice variants, and inframe frameshifts were considered VUS’s (P-score=3); the remaining variant types were considered likely benign or benign (P-score ≤2). The final P-score of a variant was the ClinVar P-score if a ClinVar annotation exists for that variant, and if not, the SnpEff P-score was used. See **Supplementary table 6** for details on pathogenicity scoring.

P-scores from pairs of mutation events (i.e. SNV, indel, or LOH) were summed to yield a biallelic pathogenicity score (BP-score), giving a maximum possible score of 10. Deep deletions were automatically given a score of 10. Per gene, the biallelic event with the highest score was taken the biallelic status of the gene. If multiple events had the same score, a biallelic event was greedily selected.

### Extracting mutation contexts

The counts of 3 types of mutation contexts (SNV, indel, and structural variant (SV) contexts) were determined from the somatic variant data from the HMF and BRCA-EU cohorts. This was performed using the R package *mutSigExtractor* (https://github.com/UMCUGenetics/mutSigExtractor).

The SNV contexts comprised of 96 trinucleotide contexts, which are composed of one of six classes of base substitutions (C>A, C>G, C>T, T>A, T>C, T>G) in combination with the immediate 5’ and 3’ flanking nucleotides.

The indel contexts comprised of 6 types based on the presence of: short tandem repeats (ins.rep, del.rep); short stretches of identical sequence at the breakpoints, also known as microhomology (ins.mh, del.mh); or the presence of neither (ins.none, del.none). Indels in repeat regions were defined as the presence of ≥1 copy of the indel sequence downstream (i.e. in the 3’ direction) from the breakpoint, where sequence length must be <50bp. Indels with flanking microhomology were defined as the presence of the following sequence features up or downstream from the breakpoint: (i) ≥1 copy of the indel sequence if the indel sequence length is ≥50bp; (ii) ≥2bp sequence identity to the indel sequence; or (iii) ≥1bp sequence identity if the indel sequence length is ≥3bp. For (ii) and (iii) the number of up or downstream bases searched was equal to the length of the indel. The 6 indel contexts types were further expanded into 30 indel contexts by stratifying ins.rep, del.rep, ins.none, and del.none by indel sequence length (1-4bp and ≥5bp); and ins.mh and del.mh by the number of bases in microhomology (‘bimh’; 1-4bp and ≥5).

The 16 SV contexts were composed of the SV type (deletion, duplication, inversion, translocation) and the SV length (1-10kb, 10-100kb, 100kb-1Mb, 1Mb-10Mb, >10Mb). Note that SV length is not applicable for translocations.

### Random forest training

#### Features

To construct the features for training the **C**lassifier of **HO**mologous **R**ecombination **D**eficiency (CHORD), the 96 trinucleotide contexts were simplified to 6 base substitution contexts by discarding the 5’ and 3’ flanking nucleotide information. For CHORD-del.mh.merged, the 30 indel contexts were simplified to the 6 indel types. For CHORD and CHORD-signature, the del.mh indel type was split into 2 bins: del.mh with 1bp homology and 2 to ≥5 (i.e. equivalent to ≥5bp) homology (del.mh.bimh.1 and del.mh.bimh.2.5 respectively). Then, relative contribution was calculated for each feature per mutation context type (i.e. SNV, indel and SV contexts separately). For CHORD-signature, the 96 trinucleotide contexts were fitted to the 30 COSMIC SBS signatures[9] using the non-negative least squares algorithm (incorporated in *mutSigExtractor*). The SV contexts were fitted in the same manner to the 6 SV signatures[9]. The relative contribution of the SBS signatures, SV signatures, and indel contexts was then calculated per mutation type.

#### Training set

The training set consisted of samples which we could confidently consider BRCA1/2 deficient or proficient based on the P-scores/BP-scores as described in *Determining gene biallelic status* and **Supplementary table 6**. BRCA1/2 deficiency was defined as having a BP-score = 10. This includes samples with: (i) a deep deletion, (ii) LOH in combination with a pathogenic SNV/indel or an out-of-frame frameshift, or (iii) two pathogenic SNV/indels and/or or out-of-frame frameshifts. Within the BRCA1/2 deficient group, samples where the absolute frequency of indels within repeat regions was >14000 were considered to have microsatellite instability (MSI) and were removed. This threshold was determined by correlating the frequency of indels within repeat regions for a selection of samples to a 5-gene PCR panel for detecting MSI (BAT25, BAT26, NR21, NR24 and MONO27 markers; data not shown). This filtering step was done as the relative contribution of indels in repeat regions are grossly overrepresented in samples with MSI, thereby masking the contribution of microhomology deletions. This sample group ultimately consisted of 35 BRCA1 (‘BRCA1’ class) and 89 BRCA2 (‘BRCA2’ class) deficient samples which were both considered HRD during the training. Conversely, BRCA1/2 proficiency required the following criteria: (i) Absence of deep deletions or LOH; (ii) all SNV/indels had a P-score ≤ 3 (VUS or lower in impact); (iii) for the highest impact pair of SNV/indels (i.e. highest BP-score), both variants had a P-score ≤ 3 (VUS or lower in impact). This BRCA proficient group (‘none’ class) consisted of 1,902 samples which were considered HRP during the training (**Supplementary figure 1**).

#### Training procedure

The training procedure for CHORD (as well as other models described in this study) is illustrated in **Supplementary figure 2**. A core training procedure, which performs feature selection and class resampling, forms the basis for the full training procedure (**Supplementary figure 2a**). Feature selection was done to retain mutation contexts which were significantly higher (p < 0.01, determined by one-tailed Wilcoxon tests) in BRCA1/2 deficient versus proficient samples. Class resampling serves to reduce the difference in the number of samples between each class (i.e. class imbalances). Here, a grid search was performed to determine the optimal pair of the following parameters: (i) down-sampling of the ‘none’ class: 1x (i.e. no down-sampling), 2x or 4x; (ii) up-sampling of the ‘BRCA1’ class: 1x (i.e. no up-sampling), 1.5x or 2x. For each iteration of the grid search, 10-fold cross-validation (CV) was performed, after which the area under the precision-recall curve (AUPRC) was calculated. The parameter pair with the highest AUPRC was chosen. With the selected features and resampling parameters, a random forest was then trained that predicts the probability of a new sample being in one of the aforementioned 3 classes (i.e. ‘BRCA1’, ‘BRCA2’ or ‘none’). We defined the HRD probability as the sum of the probability of belonging to the ‘BRCA1’ and ‘BRCA2’ classes, where a sample was considered HRD if the HRD probability was ≥0.5. Random forests were trained using the *randomForest* R package.

The full training procedure was split into 2 stages (**Supplementary figure 2b**). The first stage serves to filter ‘BRCA1’ or ‘BRCA2’ samples from the which are likely not HRD (e.g. due to reversal of biallelic inactivation via a second frameshift bringing the gene in frame), or ‘none’ samples which are likely not HRP (e.g. due to deficiencies in other HR genes). Here, the core training procedure is encapsulated by a 10-fold CV loop to allow every sample to be excluded from the training set to subsequently calculate an unbiased HRD probability. This was repeated 100 times and the number of times each sample was HRD or HRP was calculated. ‘BRCA1’ or ‘BRCA2’ samples that were predicted HRD < 60 times were blacklisted while ‘none’ samples that were predicted HRD > 40 times were blacklisted. In the second training stage, the core training procedure was performed on a training set without the blacklisted samples. This yielded the final random forest model.

The performance of the final random forest model was assessed using 2 approaches: (i) 10-fold CV of the training set by further encapsulating the full training procedure in a 10-fold CV loop; (ii) applying the final random forest model to an external dataset (BRCA-EU dataset). An AUPRC was then calculated for both approaches. In the case of the BRCA-EU dataset, BRCA1/2 deficiency annotations were obtained from Davies *et al.* 2017[9]. All performance metrics were calculated using the *mltoolkit* R package (https://github.com/UMCUGenetics/mltoolkit).

### Determining the genetic cause of HRD

To determine the genetic cause of HRD, tumors were first selected from the HMF cohort based on the absence of MSI, having ≥50 indels, and ≥30 SVs for HRD predicted samples (**Supplementary table 1**). Furthermore, for patients with multiple biopsies, a single tumor per patient was selected (based on the one with highest tumor purity). In total, 3504 tumors were selected (from the 3824 in total) to represent each patient. The following procedure was then employed for identifying biallelic loss in each of the 781 cancer/HR associated genes. First, high frequency germline SNV/indels (**Supplementary figure 15**) were marked as benign (P-score = 0). Then, each gene was screened for the following events: (i) a deep deletion; (ii) LOH in combination with a germline SNV/indel with a P-score ≥ 4 (likely pathogenic or higher in impact); (iii) LOH in combination with a somatic SNV/indel with a P-score ≥ 3 (VUS or higher in impact); or (iv) two SNVs/indels (germline + somatic, or 2x somatic) both with a P-score = 5 (pathogenic). See **Supplementary table 6** for details of the P-score thresholds used.

After applying CHORD to the HMF cohort, we then determined whether each of the 781 genes was significantly more frequently deficient in CHORD-HRD vs. CHORD-HRP patients using a one-tailed Fisher’s exact test, with multiple testing correction performed with the Hochberg procedure using the p.adjust() function in R. This was done to determine the genes most likely to cause HRD when inactivated. Six genes were found with a q-value < 0.1 and had at least 5 patients with deficiency in the corresponding gene: BRCA1, BRCA2, RAD51C, PALB2, NF1, and STARD13 (**Supplementary figure 16**). NF1 and STARD13 have not been reported to be involved in HR, and thus further analyses were performed to validate the enrichment for these 2 genes.

Since BRCA1 and NF1 are both located on Chr17, we reasoned that copy number alterations (CNA; in this case referring to deep deletions or LOH) that affect BRCA1 also affect NF1. This leads to frequent biallelic loss of NF1 even though the gene is likely not associated with HRD. A similar situation was suspected for BRCA2 and STARD13 which are both located on Chr13. Thus, one-tailed Fisher’s exact tests were performed to determine whether CNAs in each of the 781 genes significantly co-occurred more often with a CNA in BRCA1 or BRCA2. Multiple testing correction was performed using the Hochberg procedure. Indeed, enrichment in the co-occurrence of BRCA1 and NF1 CNAs was found, and was similarly the case for BRCA2 and STARD13 (**Supplementary figure 17**). We thus concluded that biallelic loss of NF1 and STARD13 are likely not associated with HRD and were therefore excluded from **Figure 3a**.

### Clustering of CHORD-HRD samples

Clustering of CHORD-HRD samples based on biallelic inactivating events (as in **Figure 3c**) is illustrated in **Supplementary figure 22**. First, samples were split into 4 groups according to their HRD subtype and whether a sample had an impactful biallelic event (P-score pair of 5 and ≥3).

For each of these groups, the HRD associated gene with the max BP-score was greedily determined per sample and assigned a score of 1, with the remaining genes being assigned a score of 0. Genes were prioritized as follows BRCA2, BRCA1, RAD51C, PALB2. This was based on highest to lowest enrichment of gene deficiency in CHORD-HRD vs. CHORD-HRP as described above. With the resultant (1,0) matrix, a sorting operation was performed. A post-processing step (done purely for cosmetic purposes) ensured that samples with deep deletions, LOH + frameshift, and LOH + other SNV/indels in the corresponding gene were ranked first. The sorted (1,0) matrices from the 4 sample groups were combined, and consecutive rows of 1’s were considered a cluster. For the 2 groups representing samples with no impactful biallelic event, all samples were considered to be in one cluster. These 2 groups corresponded to clusters 4 and 6 in **Figure 3c**, and samples in these clusters were considered to have an unknown cause of HRD.

For **Supplementary figure 20**, samples were first split by cancer type before performing the above procedure.

### Code availability

CHORD is available as an R package at https://github.com/UMCUGenetics/CHORD.

### Data availability

WGS data and corresponding metadata have been obtained from the Hartwig Medical Foundation and provided under data request numbers DR-010 and DR-047. Both WGS data and metadata is freely available for academic use from the Hartwig Medical Foundation through standardized procedures and request forms can be found at https://www.hartwigmedicalfoundation.nl.

## Supplementary tables

**Supplementary table 1**: Predictions from CHORD and CHORD-signature on the HMF, BRCA-EU, and PCAWG datasets as well as metadata for each sample

**Supplementary table 2**: Mutation contexts and mutational signatures extracted from the HMF, BRCA-EU, and PCAWG datasets

**Supplementary table 3**: List of 781 cancer and HR related genes used for the pan-cancer analysis of HRD and results of the enrichment analysis to determine HRD associated genes

**Supplementary table 4**: Genotypes of BRCA1/2, RAD51C, PALB2 and other HR genes for the 310 CHORD-HRD patients from the HMF and PCAWG datasets

**Supplementary table 5**: Incidence and genetic cause of HRD by cancer type in the HMF and PCAWG dataset

**Supplementary table 6**: Pathogenicity scoring of variants used to determine biallelic gene status, including biallelic pathogenicity score inclusion criteria for CHORD training data

## Supplementary information

### 1. Prerequisites for accurate HRD prediction

It is important to note that microsatellite instability (MSI) negatively affects CHORD’s ability to accurately predict HRD. MSI is a hypermutator phenotype characterized by an exceptional number of indels in regions with (tandem) repeats. As CHORD uses relative values of mutation contexts as input, MSI (defined as having more than 14,000 indels within repeat regions) results in a reduction of the relative contribution of microhomology deletions and thus an underestimated HRD probability (‘false negatives’). The figure below shows the CHORD being applied to all 3,824 HMF samples. All MSI samples were predicted HRP by CHORD even though 4 of these samples had biallelic loss of BRCA2. This could be circumvented by incorporating a MSI trained HRD classifier within CHORD. However, the number of HRD predicted samples with MSI is currently too small for training such a classifier and therefore we have built in MSI status checking as a quality control (QC) step within CHORD.

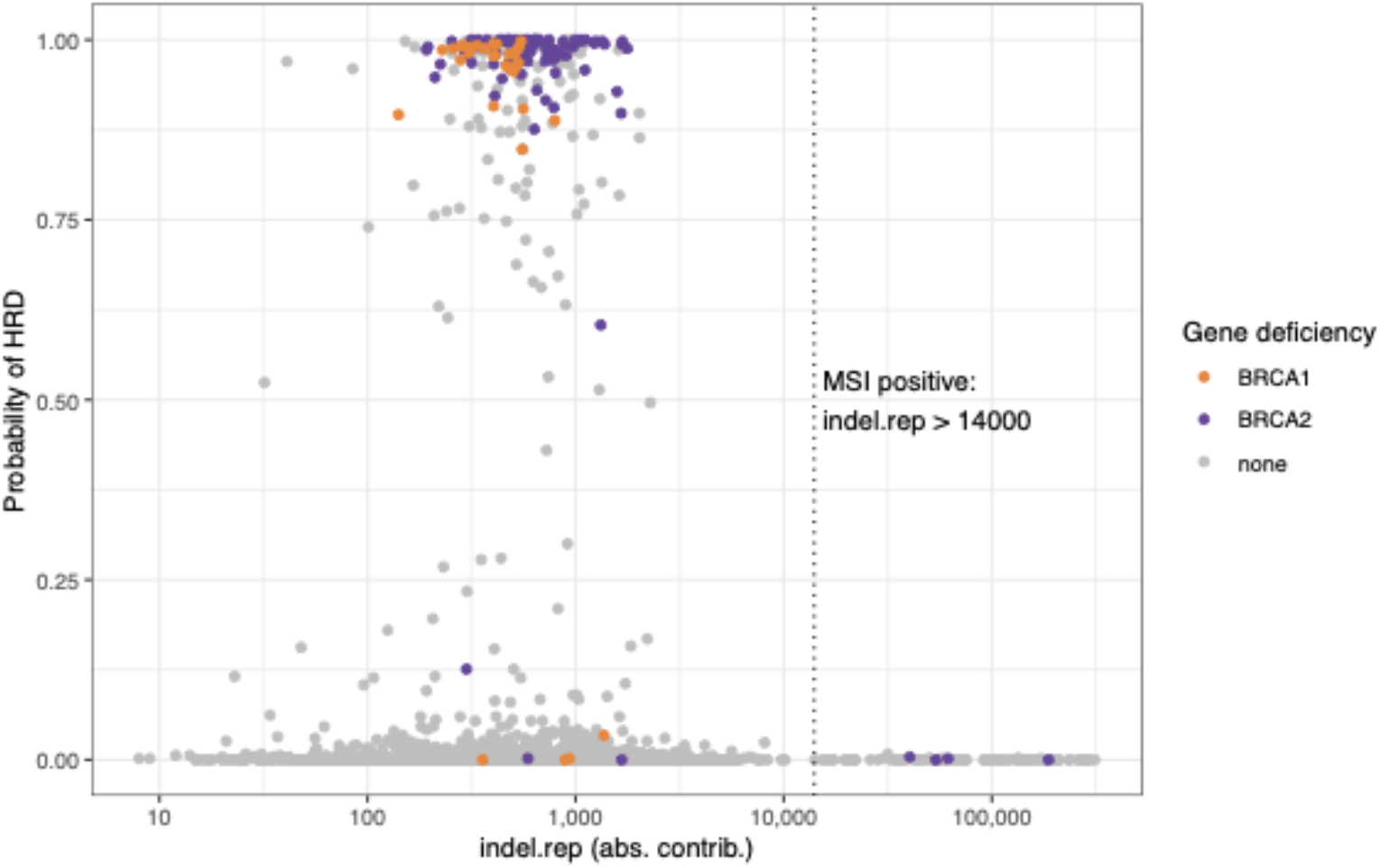

A minimum number of mutations is also required for accurate HRD prediction. To determine this, we progressively down-sampled all mutations for each sample in the training set and measured the reduction in performance. We found that at least 50 indels were required for accurately predicting HRD, and if a sample was predicted HRD, at least 30 SVs were required for distinguishing BRCA1-type from BRCA2-type HRD. These threshold levels may be particularly relevant for samples with low tumor purity and/or read coverage, and are as such also included as a QC step within CHORD.

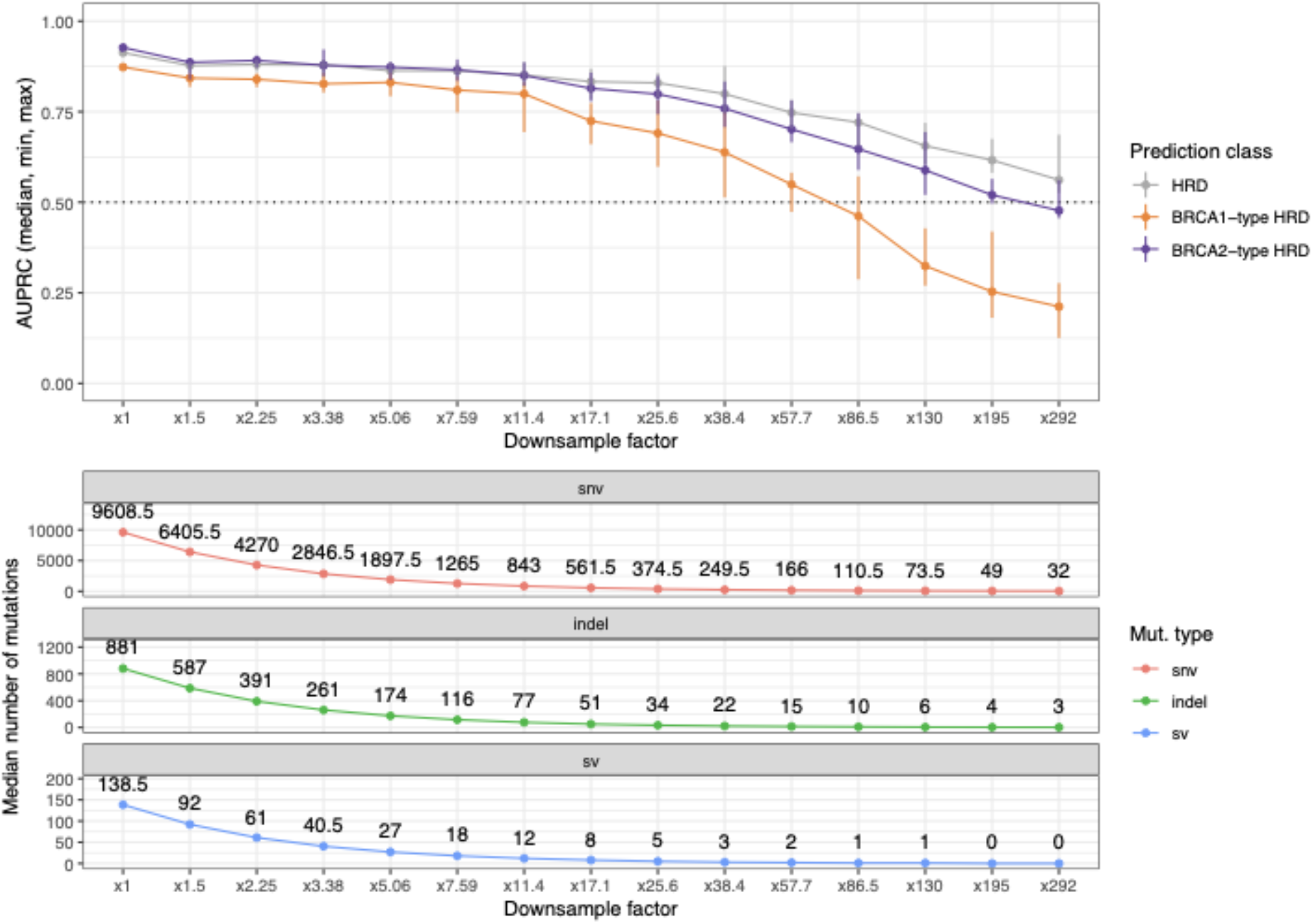

### 2. CHORD as a tool to uncover novel pathogenic variants

The HRD predictions from CHORD can provide supportive evidence for interpreting variants of unknown significance (VUS), either germline or somatic, which can in turn be used for prioritizing rare VUS’s for experimental validation as shown in the scheme below.

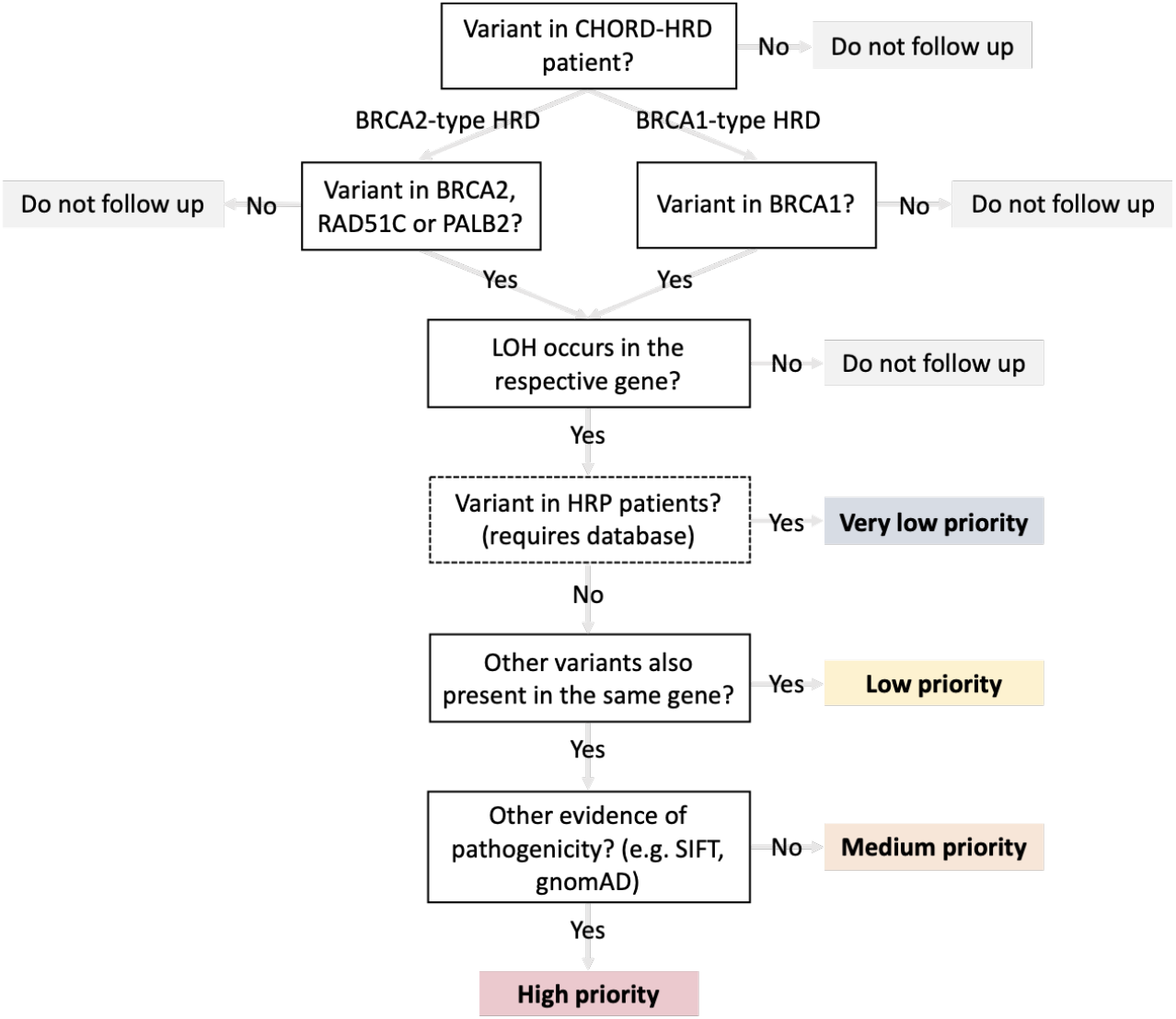

From 13 CHORD-HRD patients, we identified 14 variants not previously described to be pathogenic but which in combination with LOH could explain biallelic loss of a HR gene. Moreover, biallelic loss of the respective gene corresponded to the associated HRD subtype, and all of these variants were not present in HRP samples, providing additional support that these variants are potentially pathogenic. From these variants, 2 *BRCA2* missense variants (c.9230T>C, c.9254C>T) were both found in patient HMF000429. Here, further validation is required to determine which was the driver mutation (or possibly the combination). The remaining 12 variants were the sole variant found in the respective gene and respective patient, and are thus more likely to be pathogenic driver variants. Of these, one *BRCA2* missense variant (c.8045C>T) had a ‘Uncertain_significance’ annotation in ClinVar but a low population frequency according to gnomAD as well as being predicted as ‘deleterious’ by SIFT and ‘probably damaging’ by PolyPhen, supporting its potential pathogenicity.

| sample_index | cluster | sample | chrom | pos | ref | alt | variant_type | mutation_origin | hgnc_symbol | clinvar_annotation | gnomad_annotation | manual_interpretation |
| --- | --- | --- | --- | --- | --- | --- | --- | --- | --- | --- | --- | --- |
| 109 | 1 | efbec43c-0c16-4006-abe8-c3ec2ec42c05 | 13 | 32912849 | A | T | stop_gained | somatic | BRCA2 |  |  | likely_pathogenic |
| 110 | 1 | HMF002507A | 13 | 32907215 | G | T | stop_gained | somatic | BRCA2 |  |  | likely_pathogenic |
| 111 | 1 | HMF002917A | 13 | 32914182 | T | A | stop_gained | germline | BRCA2 |  |  | likely_pathogenic |
| 112 | 1 | 53534b3c-cd15-4d68-a9b1-6902bb234c45 | 13 | 32944694 | G | C | missense_variant | somatic | BRCA2 |  |  | likely_pathogenic |
| 113 | 1 | HMF000429B | 13 | 32954256 | T | C | missense_variant | somatic | BRCA2 |  |  | VUS |
| 113 | 1 | HMF000429B | 13 | 32954280 | C | T | missense_variant | somatic | BRCA2 |  |  | VUS |
| 114 | 1 | HMF002161A | 13 | 32937384 | C | T | missense_variant | somatic | BRCA2 | Uncertain_significance | AF=0.00001194;polyphen=probably_damaging;SIFT=deleterious | pathogenic |
| 115 | 1 | HMF003376A | 13 | 32954283 | G | T | splice_donor_variant | somatic | BRCA2 |  |  | likely_pathogenic |
| 116 | 1 | ad007b59-46f6-48b9-958a-2b9d28cca3b7 | 13 | 32900222 | CTTTGT | C | intron_variant | germline | BRCA2 | Uncertain_significance |  | VUS |
| 117 | 1 | HMF002299A | 13 | 32931943 | A | C | missense_variant | germline | BRCA2 |  |  | likely_pathogenic |
| 122 | 2 | HMF000801A | 17 | 56780605 | A | C | missense_variant | somatic | RAD51C |  |  | likely_pathogenic |
| 123 | 2 | HMF003585A | 17 | 56774098 | GA | TG | missense_variant | somatic | RAD51C |  |  | likely_pathogenic |
| 245 | 5 | cf6c7066-6611-43ec-aef1-295a976cce5b | 17 | 41276094 | C | A | missense_variant | somatic | BRCA1 |  |  | likely_pathogenic |
| 246 | 5 | HMF002751A | 17 | 41245270 | T | A | missense_variant | somatic | BRCA1 |  |  | likely_pathogenic |

### 3. CHORD can detect HRD in a substantial number of cases that would be missed by genetic testing

To assess the potential value of CHORD in a clinical setting, we compared CHORD’s predictions for HMF patients to the hypothetical outcomes of common genetic testing approaches. In the clinic, HRD detection is currently often done by screening for pathogenic BRCA1/2 SNVs/indels based on annotations from curated databases (e.g. ClinVar). This is performed either on blood biopsies (blood genetic testing), analogous to screening for germline SNVs/indels in sequencing data of the HMF cohort (which was analysed by tumor-normal pair whole genome sequencing); or on tumor biopsies (tumor genetic testing), analogous to screening both germline and somatic SNVs/indels. Our genetic analyses (below figure) indicate that blood genetic testing would identify a pathogenic BRCA1/2 SNV/indel (according to ClinVar; or an out-of-frame frameshift) in 18% of CHORD-HRD breast/ovarian cancer patients (which genetic testing is often restricted to), while tumor genetic testing would increase this proportion to 25%. If patients with other cancer types would be included, blood and tumor genetic testing would identify 31% and 42% of CHORD-HRD tumors (respectively) with a pathogenic BRCA1/2 SNV/indel (**a: columns 1 and 2**).

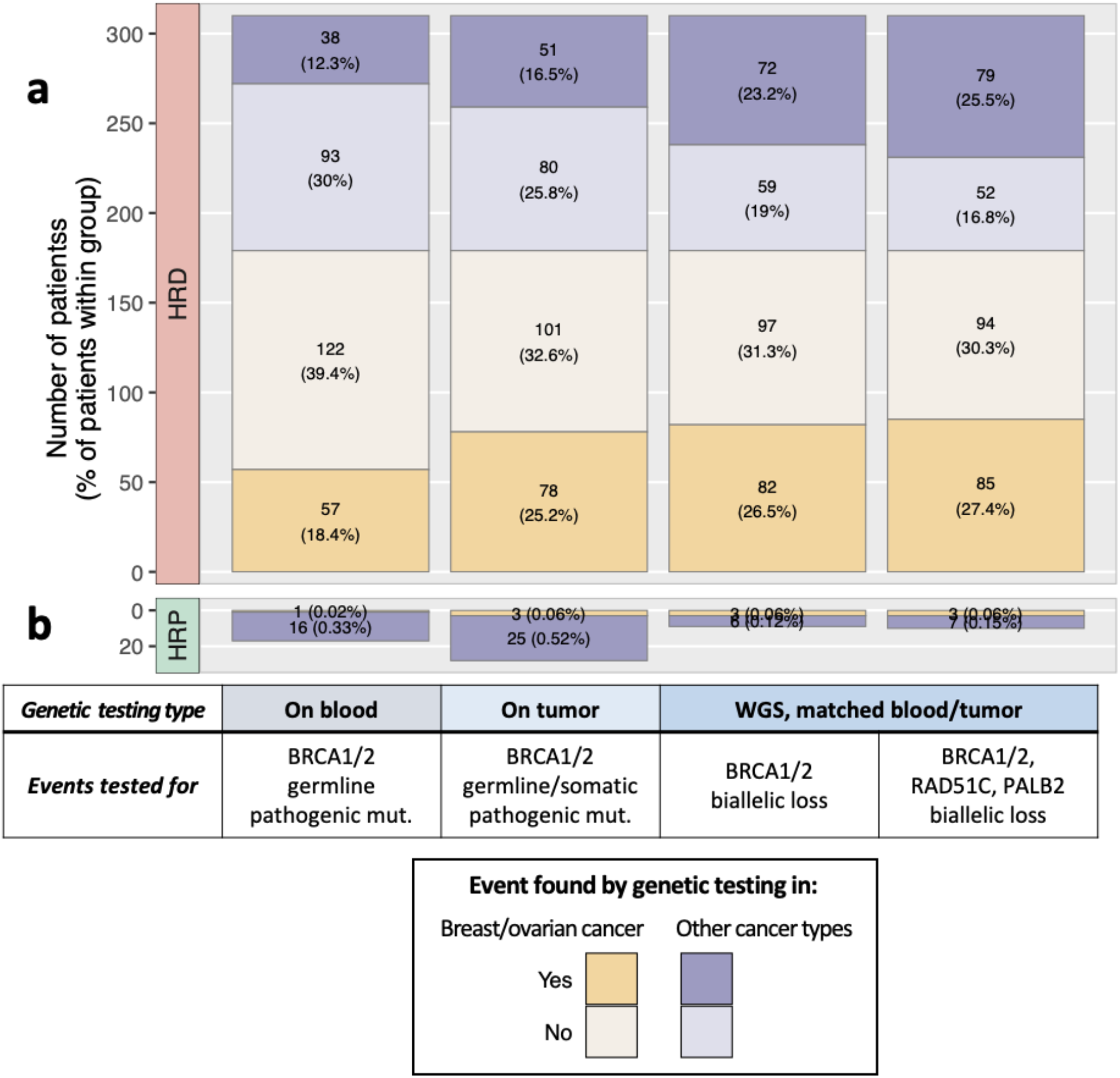

CHORD identifies a large proportion of HRD patients that would be missed by genetic testing. (**a**) and (**b**) show the percentage of CHORD-HRD or CHORD-HRP patients (respectively) from the HMF dataset in which a pathogenic event was found based on four genetic testing setups. In the first two setups (‘On blood’, ‘On tumor’), a pathogenic event was identified if a pathogenic SNV/indel was found on one allele, which was defined as a frameshift, or a likely pathogenic or pathogenic variant according to ClinVar. In the ‘WGS based testing’ setups, the pathogenic SNV/indel must also have occurred in combination with loss of heterozygosity (LOH); or, a deep deletion was identified. Only data from samples that passed CHORD’s QC criteria are shown in this figure (MSI absent, ≥50 indels, and ≥30 SVs if a sample was predicted HRD).

While not currently routinely performed in the clinic, WGS based genetic testing with matched blood/tumor biopsies (WGS genetic testing) would allow the detection of any event (including structural events such as LOH or deep deletions) that contributes to HR gene inactivation, and enables the determination of biallelic gene status. Our analyses show that WGS genetic testing would increase the number of patients that are considered HRD compared to SNV/indel-based blood/tumor genetic testing, with biallelic loss of BRCA1/2 being identified in 27% of CHORD-HRD breast/ovarian cancer patients and in 50% of patients pan-cancer (**a: column 3**). Additionally, WGS genetic testing would consider 9 CHORD-HRP patients as HRD (**b: column 3**; ‘genetic testing false positives’; these could also be CHORD false negatives), a marked decrease compared to tumor genetic testing which would identify 28 false positives (**b: column 2**). By including the two other main HRD associated genes (RAD51C and PALB2) in WGS genetic testing, biallelic gene loss would be identified in 53% of patients (**a: column 4**), while only increasing the number of genetic testing false positives from 9 to 10 patients (**b: column 5**). Our findings show that while WGS genetic testing (for biallelic loss) offers improved detection of HRD patients compared to testing for pathogenic SNVs/indels, it still misses around half of HRD patients as classified by CHORD.

## Supplementary figures

**Supplementary figure 1:**
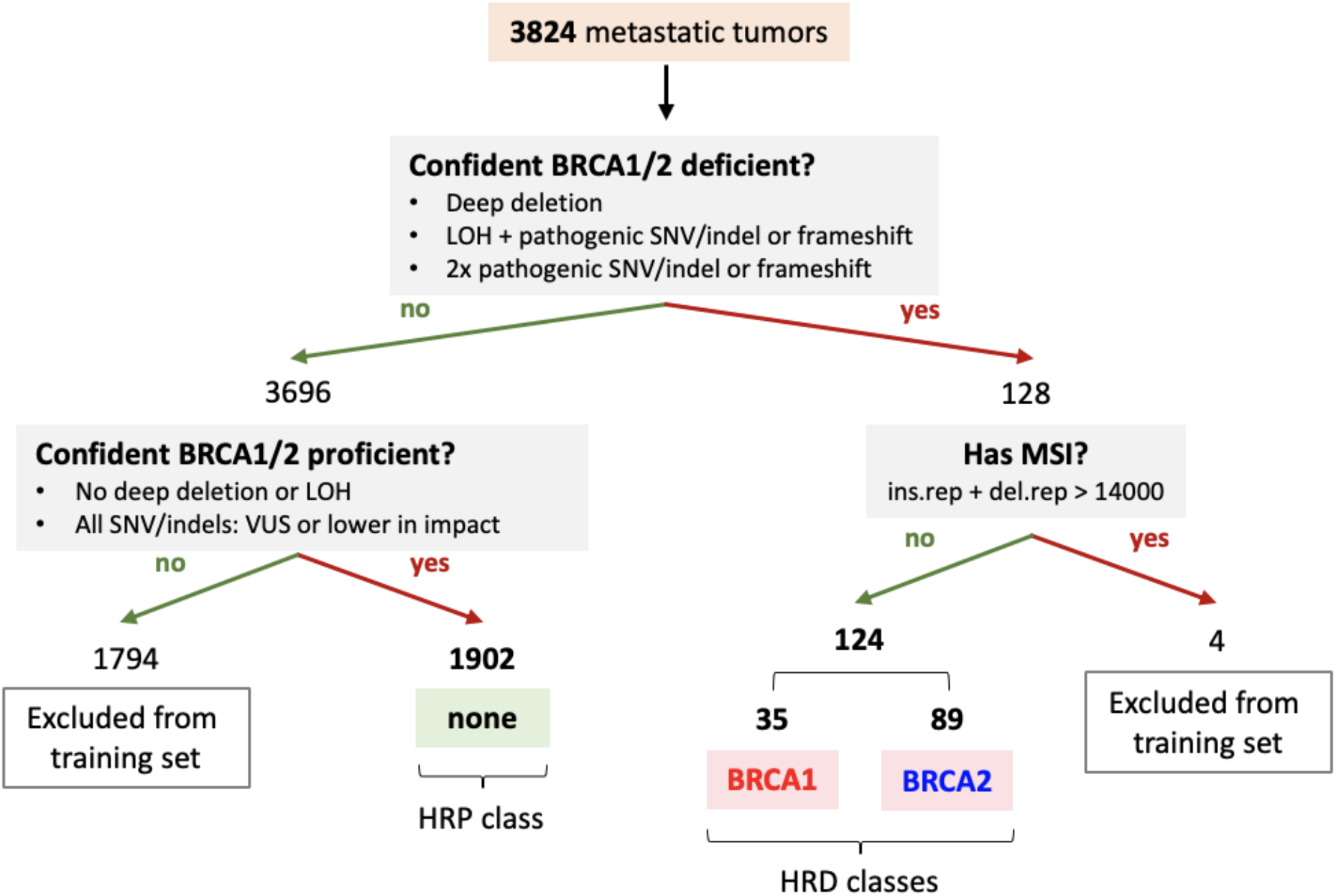
Sample selection for training CHORD. Only samples where biallelic inactivation of BRCA1/2 could confidently be identified (and did not have MSI), or where no disruption of BRCA1/2 was apparent were recruited into the training set. A total of 2,026 samples were eventually selected, consisting of 35 BRCA1 and 89 BRCA2 deficient which were both considered HRD during training, and 1,902 BRCA1/2 proficient (‘none’) which were considered HRP.

**Supplementary figure 2:**
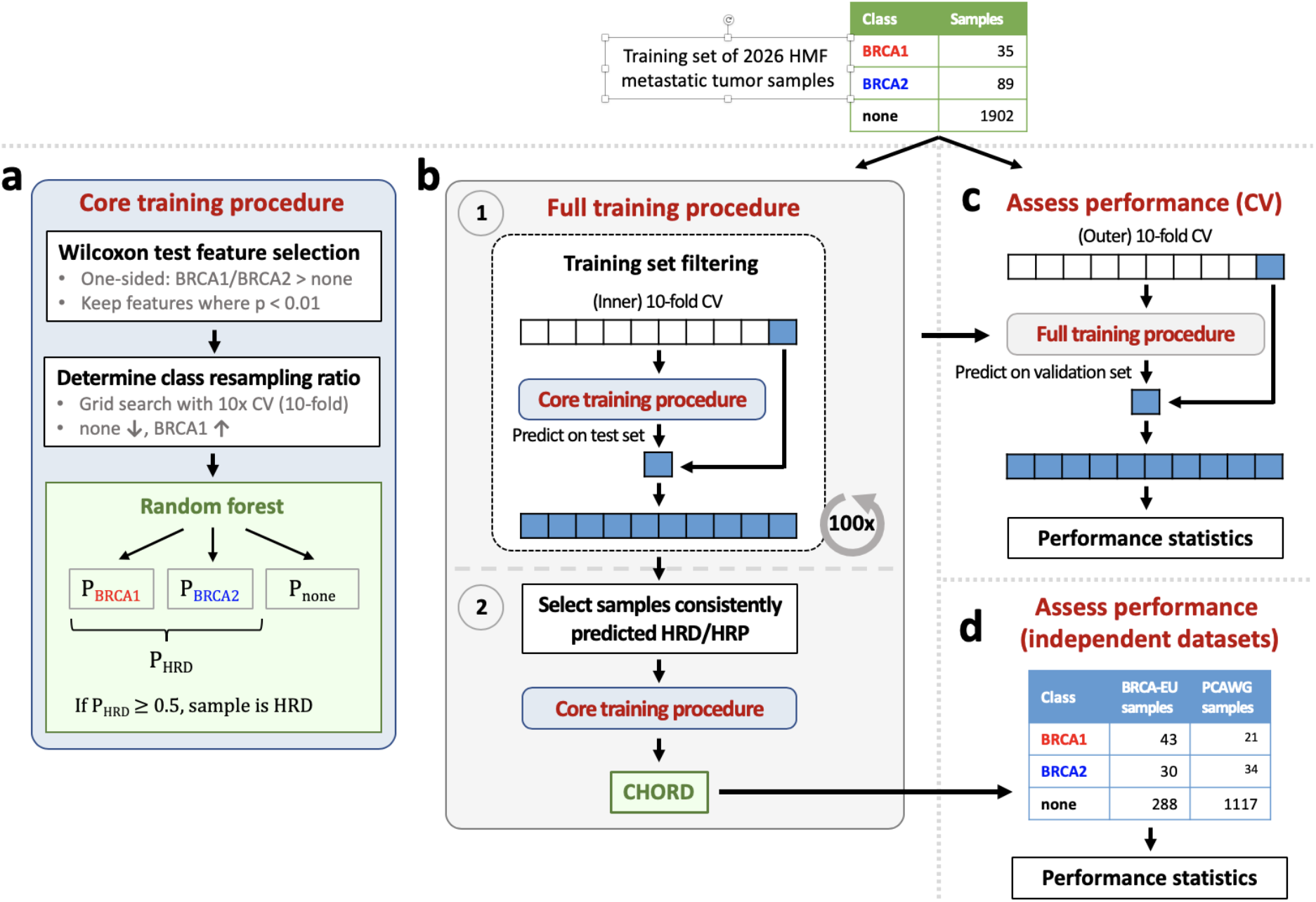
Workflow for training CHORD. ‘BRCA1’, ‘BRCA2’ and ‘none’ classes refer to BRCA1 deficient, BRCA2 deficient and BRCA1/2 proficient sample groups, respectively. (**a**) The core training procedure on which the full training procedure is based on performs feature selection and class resampling. This returns a random forest that outputs the probability of a new sample being in one of the aforementioned 3 classes. The probability of HRD (P_HRD_) is the sum of the probability of belonging to the ‘BRCA1’ and ‘BRCA2’ classes, where a sample is considered HRD if P_HRD_ is ≥ 0.5. (**b**) The full training procedure is split into 2 stages. The first stage serves to blacklist ‘BRCA1’ or ‘BRCA2’ class samples which are likely not HRD (e.g. due to reversal of biallelic inactivation via a secondary frameshift), or ‘none’ samples which are likely not HRP (e.g. due to deficiencies in other HR genes). Here, the core training procedure is encapsulated by a 10-fold cross-validation (CV) loop to allow every sample to be excluded from the training set to subsequently calculate an unbiased P_HRD_. This was repeated 100 times and the number of times each sample was HRD or HRP was calculated. ‘BRCA1’ or ‘BRCA2’ samples that were predicted HRD < 60 times were blacklisted. ‘none’ samples that were predicted HRD > 40 times were blacklisted. In the second training stage, the core training procedure was performed on a training set without the blacklisted samples. This produced the final random forest model, CHORD. (**c**) The full training procedure was further encapsulated by a 10-fold CV to assess the performance of CHORD. (**d**) Performance was also assessed by applying CHORD on two independent datasets: BRCA-EU (371 primary breast tumors) and PCAWG (1176 primary tumors, pan-cancer).

**Supplementary figure 3:**
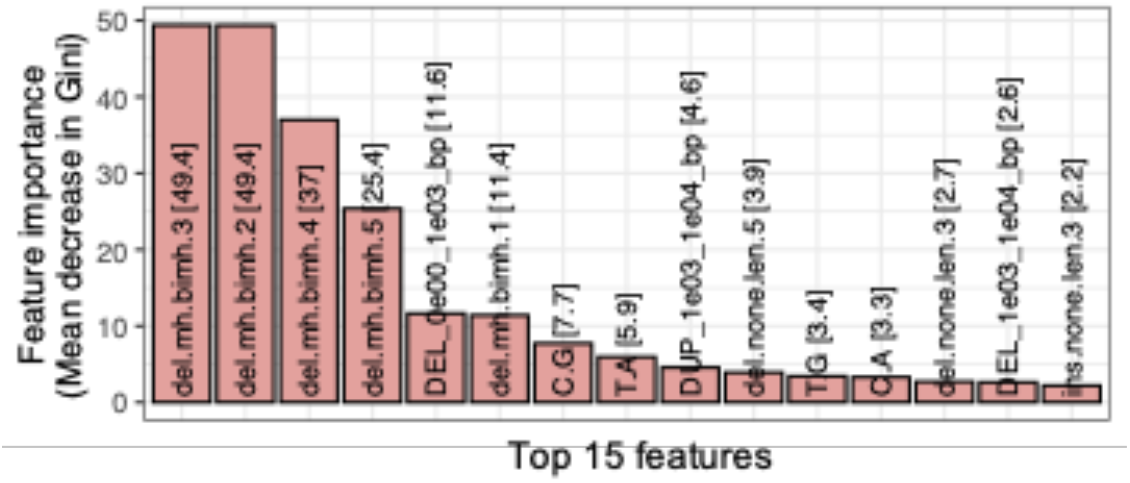
A random forest was applied to the training set revealed that deletions with ≥2bp flanking homology (del.mh.bimh.5 represents ≥5bp flanking homology) were the most important features for predicting BRCA1/2 deficiency.

**Supplementary figure 4:**
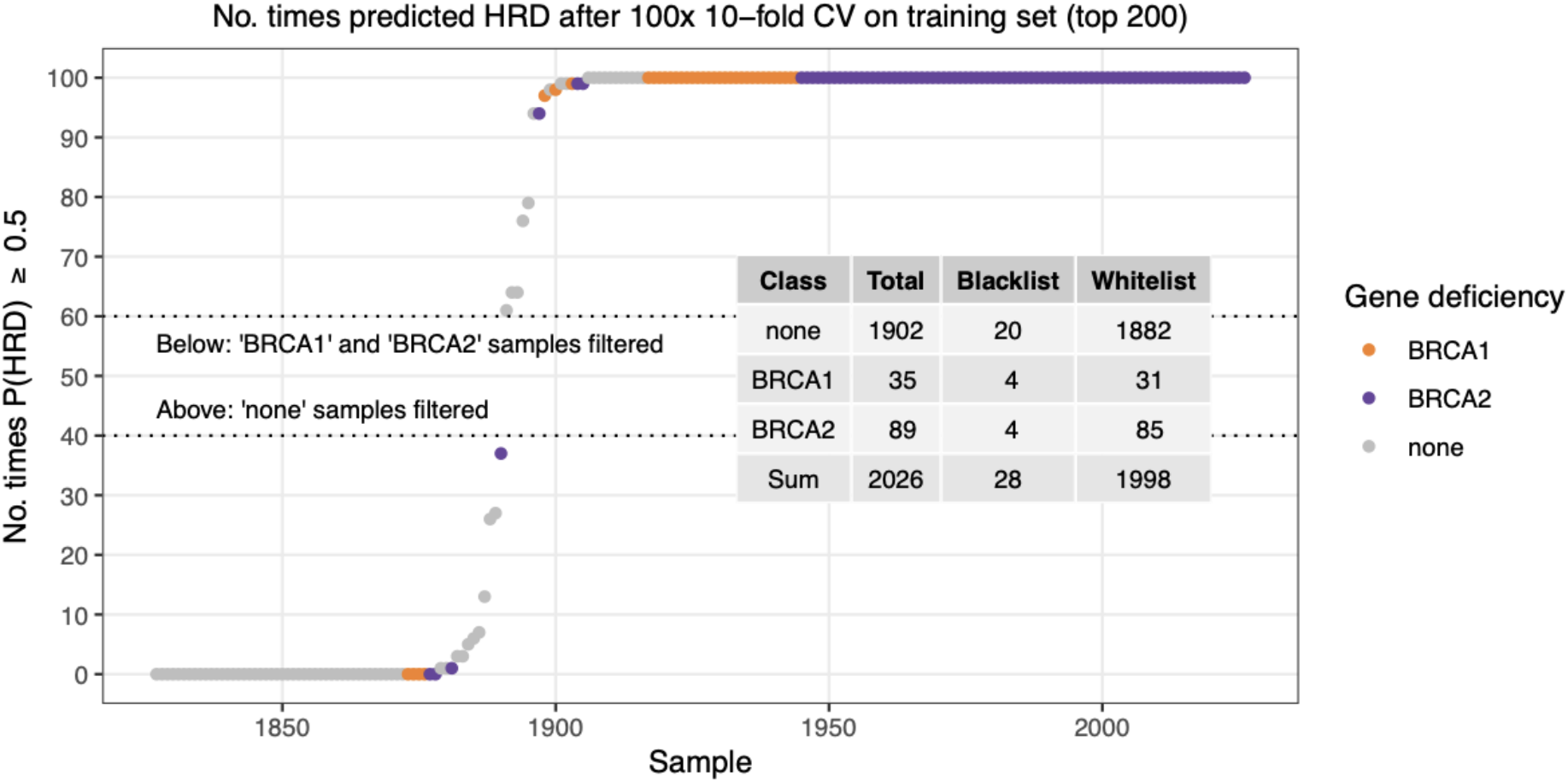
Results of the 100x repeated 10-fold CV procedure performed when training CHORD (see **Supplementary figure 2b** for details) in order to blacklist ‘BRCA1’ or ‘BRCA2’ class samples that were consistently predicted HRP (no. times HRD < 60), or ‘none’ class samples that were consistently predicted HRD (no. times HRD > 40). Samples with a probability of HRD ≥ 0.5 were considered HRD. The overlaid table summarizes the number of samples per class before and after removing the blacklisted samples from the training set. Note that only the top 200 samples are shown here as the remaining samples belonged to the ‘none’ class and were always predicted HRP.

**Supplementary figure 5:**
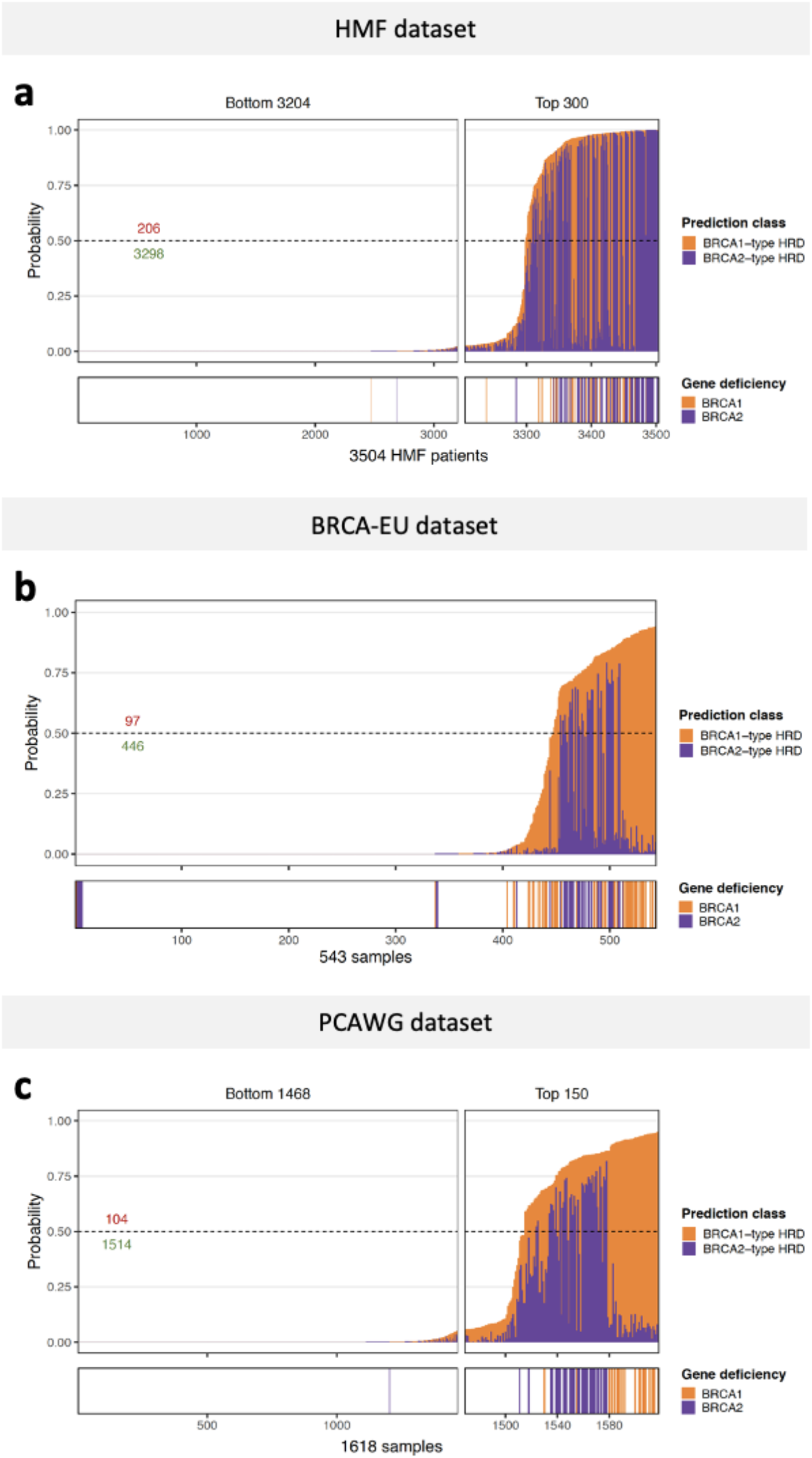
CHORD predictions on all (**a**) HMF, (**b**) BRCA-EU and (**c**) PCAWG patients. The probability of HRD for each sample (total bar height) with each bar being divided into segments indicating the probability of BRCA1- (orange) and BRCA2-type HRD (purple). Stripes below the bar plot indicate biallelic loss of BRCA1 or BRCA2. Only samples that passed CHORD’s QC criteria are shown (MSI negative, ≥50 indels, and ≥30 SVs if a sample was predicted HRD)

**Supplementary figure 6:**
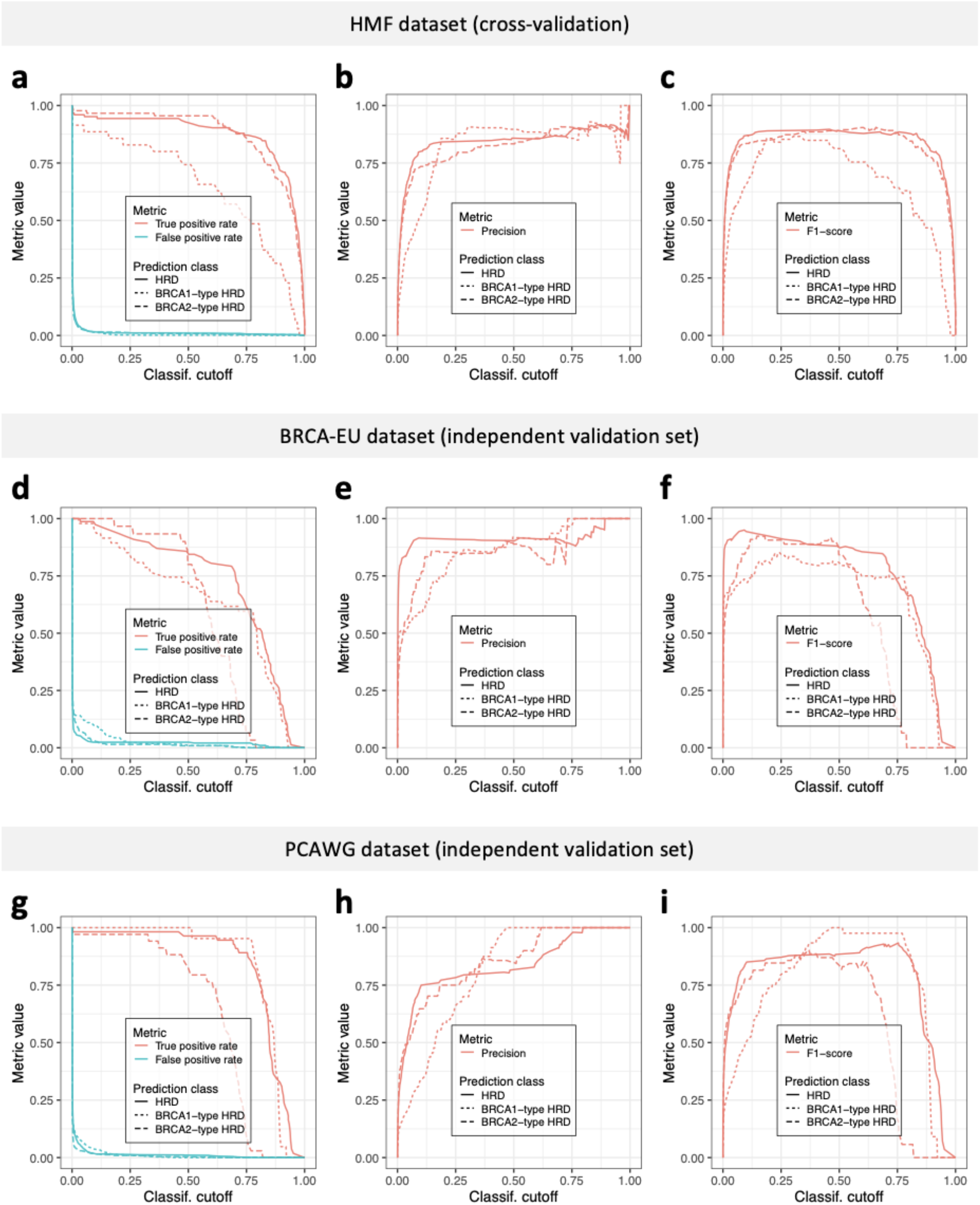
Additional performance metrics for CHORD as determined by 10-fold cross-validation (CV) on the HMF training data (**a-c**) or prediction on two independent datasets, BRCA-EU (primary breast cancer dataset; **d-f**) and PCAWG (primary pan-cancer dataset; **g-i**). Data from the BRCA-EU and PCAWG datasets are from samples that passed CHORD’s QC criteria (i.e. MSI absent, ≥50 indels, ≥30 SVs if a sample was predicted HRD).

**Supplementary figure 7:**
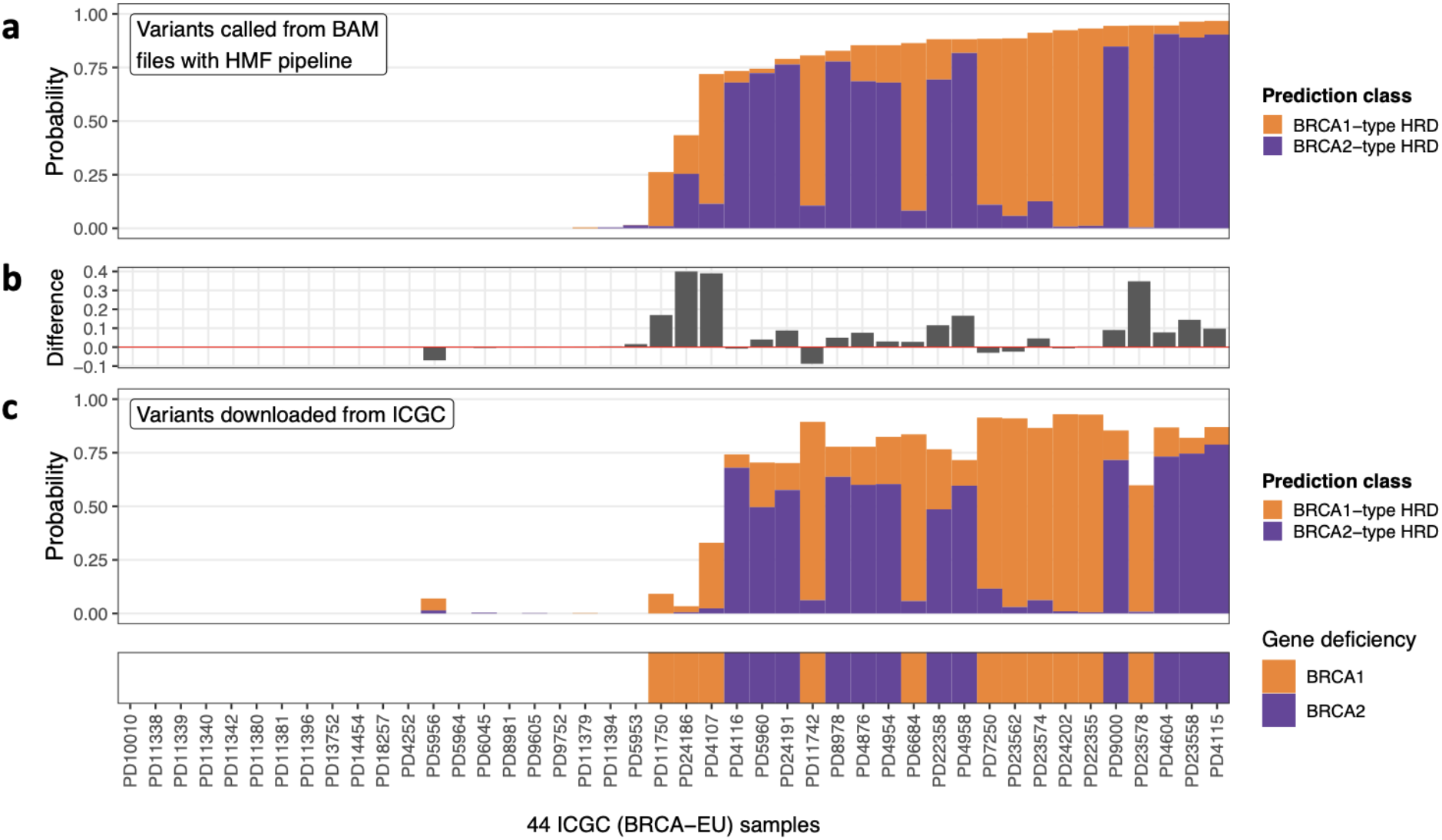
Variant calling pipeline differences affects CHORD performance. (**a**) Using variants called with the native pipeline of the HMF dataset (HMF pipeline) for HRD prediction with CHORD resulted in overall higher HRD probabilities in BRCA1/2 deficient tumors when compared to (**c**) using variants downloaded from ICGC. The differences in HRD probabilities are quantified in (**b**). All samples shown here passed CHORD’s QC criteria (i.e. MSI absent, ≥50 indels, ≥30 SVs if a sample was predicted HRD)

**Supplementary figure 8:**
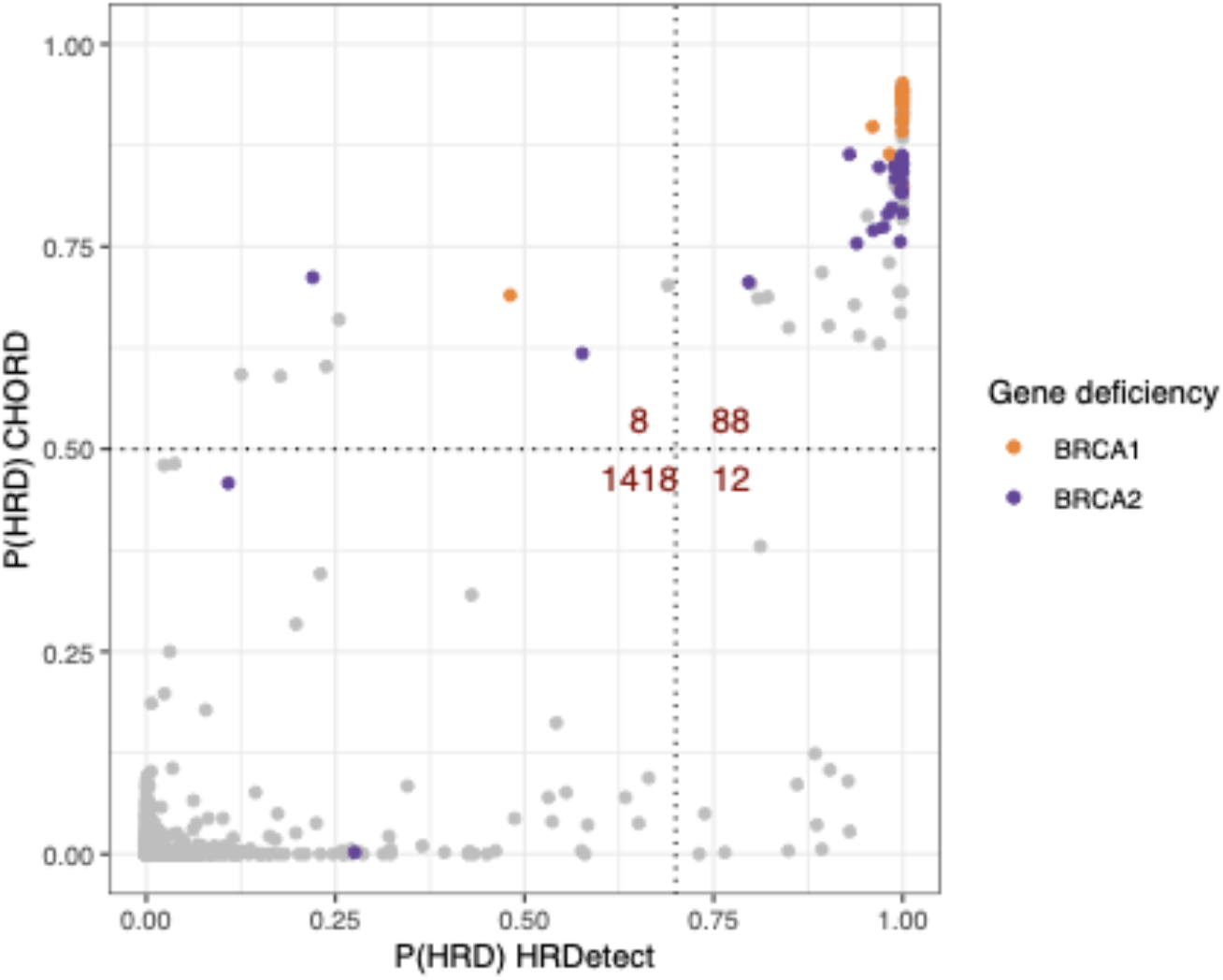
CHORD vs HRDetect predictions on the PCAWG dataset. The 1526 samples shown here had predictions from HRDetect and passed CHORD’s QC criteria for predicting HRD (i.e. MSI absent, ≥50 indels, ≥30 SVs if a sample was predicted HRD). The dotted lines indicate the classification thresholds for the two models (CHORD: 0.5, HRDetect: 0.7).

**Supplementary figure 9:**
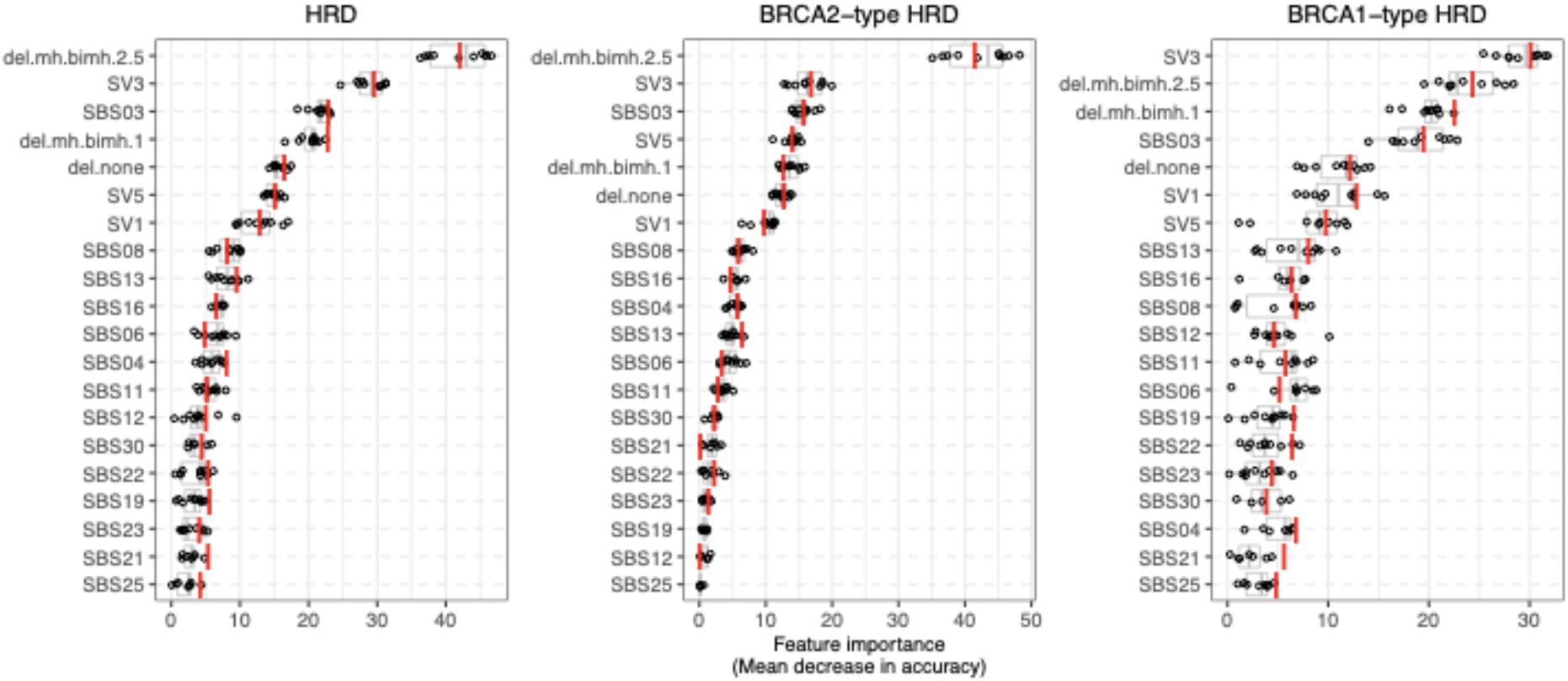
The features used by CHORD-signature to predict HRD as well as BRCA1-type HRD and BRCA2-type HRD, with their importance indicated by mean decrease in accuracy. SV# refers to the 6 SV mutational signatures and SBS# refers to the 30 single base substitution signatures as used by HRDetect. Boxplot and dots show the feature importance over 10-folds of nested CV on the training set, with the red line showing the feature importance in the final CHORD model. Boxes show the interquartile range (IQR) and whiskers show the largest/smallest values within 1.5 times the IQR.

**Supplementary figure 10:**
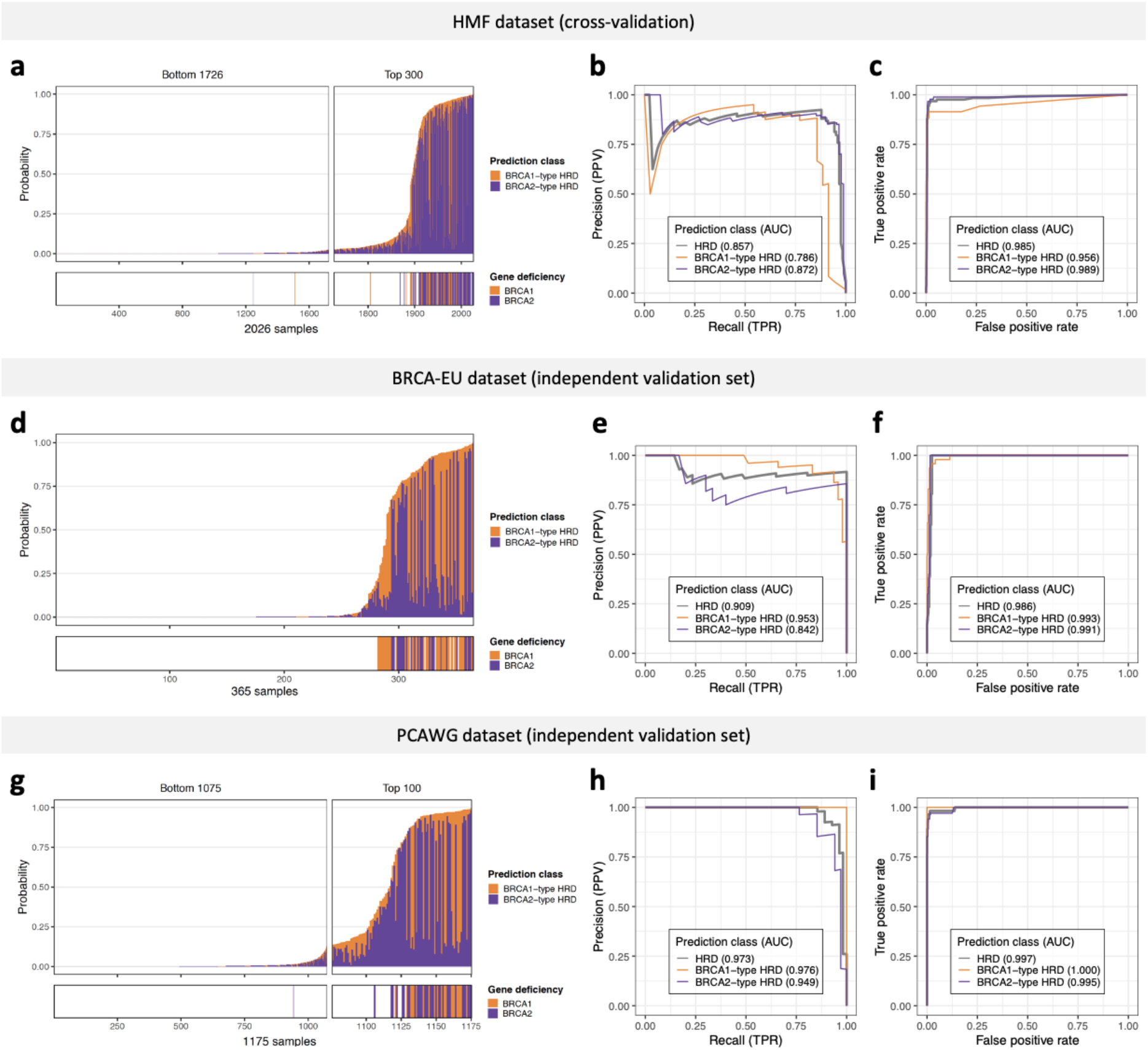
Performance of CHORD-signature as determined by 10-fold cross-validation (CV) on the HMF training data or prediction on two independent datasets: BRCA-EU (primary breast cancer dataset) and PCAWG (primary pan-cancer dataset). BRCA-EU and PCAWG samples shown here all passed CHORD’s QC criteria (i.e. MSI absent, ≥50 indels, ≥30 SVs if a sample was predicted HRD) (**a, d, g**) The probability of HRD for each sample (total bar height) with each bar being divided into segments indicating the probability of BRCA1-(orange) and BRCA2-type HRD (purple). Stripes below the bar plot indicate biallelic loss of BRCA1 or BRCA2. In (**a**), probabilities have been aggregated from the 10 CV folds. (**b, e, h**) Receiver operating characteristic (ROC) and (**c, f, i**) precision-recall curves (PR) and respective area under the curve (AUC) values showing the performance of CHORD when predicting HRD as a whole (grey), BRCA1-type HRD (orange), or BRCA2-type HRD (purple).

**Supplementary figure 11:**
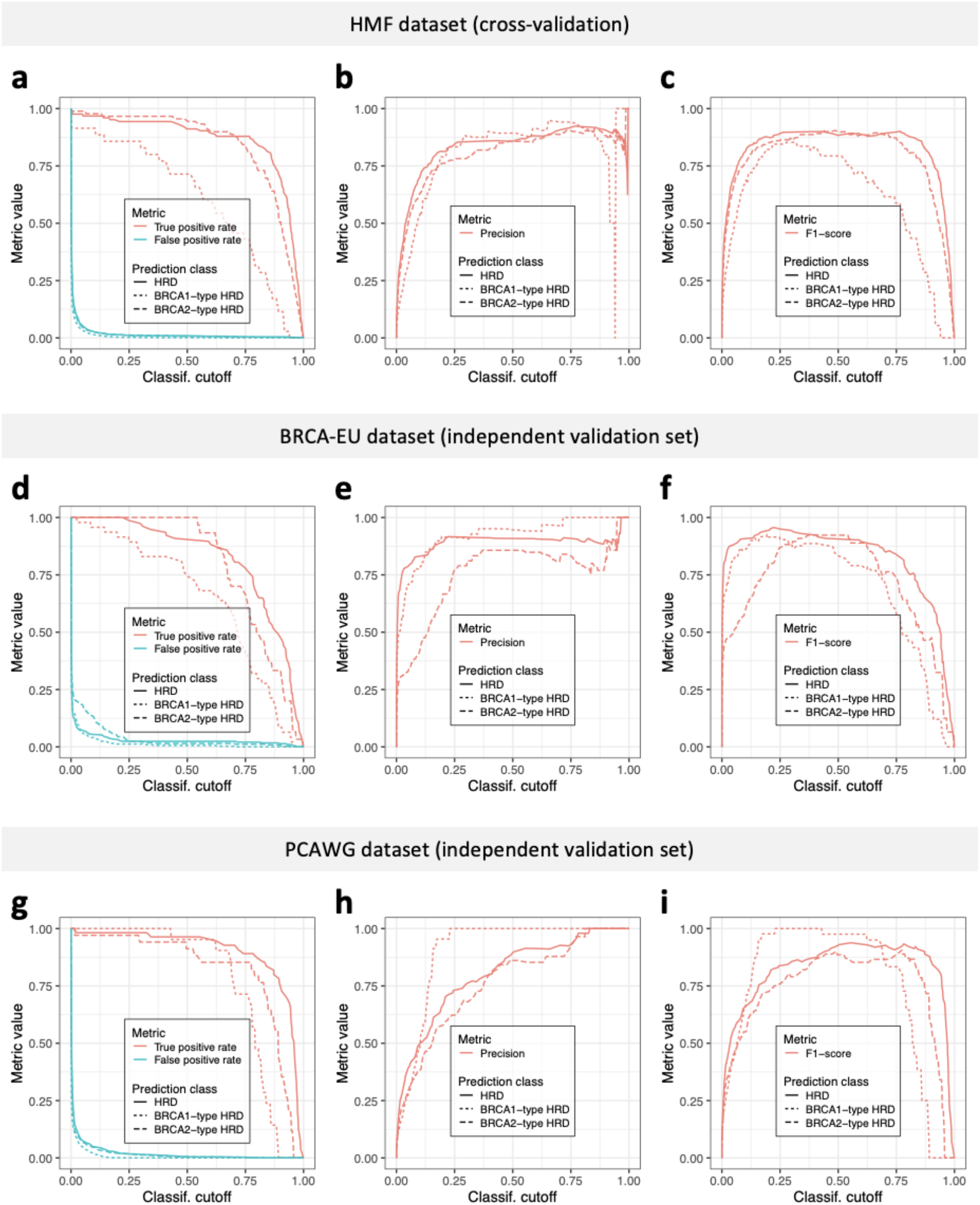
Additional performance metrics for CHORD-signature as determined by 10-fold cross-validation (CV) on the HMF training data (**a-c**) or prediction on two independent datasets, BRCA-EU (primary breast cancer dataset; **d-f**) and PCAWG (primary pan-cancer dataset; **g-i**). Data from the BRCA-EU and PCAWG datasets are from samples that passed CHORD’s QC criteria (i.e. MSI absent, ≥50 indels, ≥30 SVs if a sample was predicted HRD).

**Supplementary figure 12:**
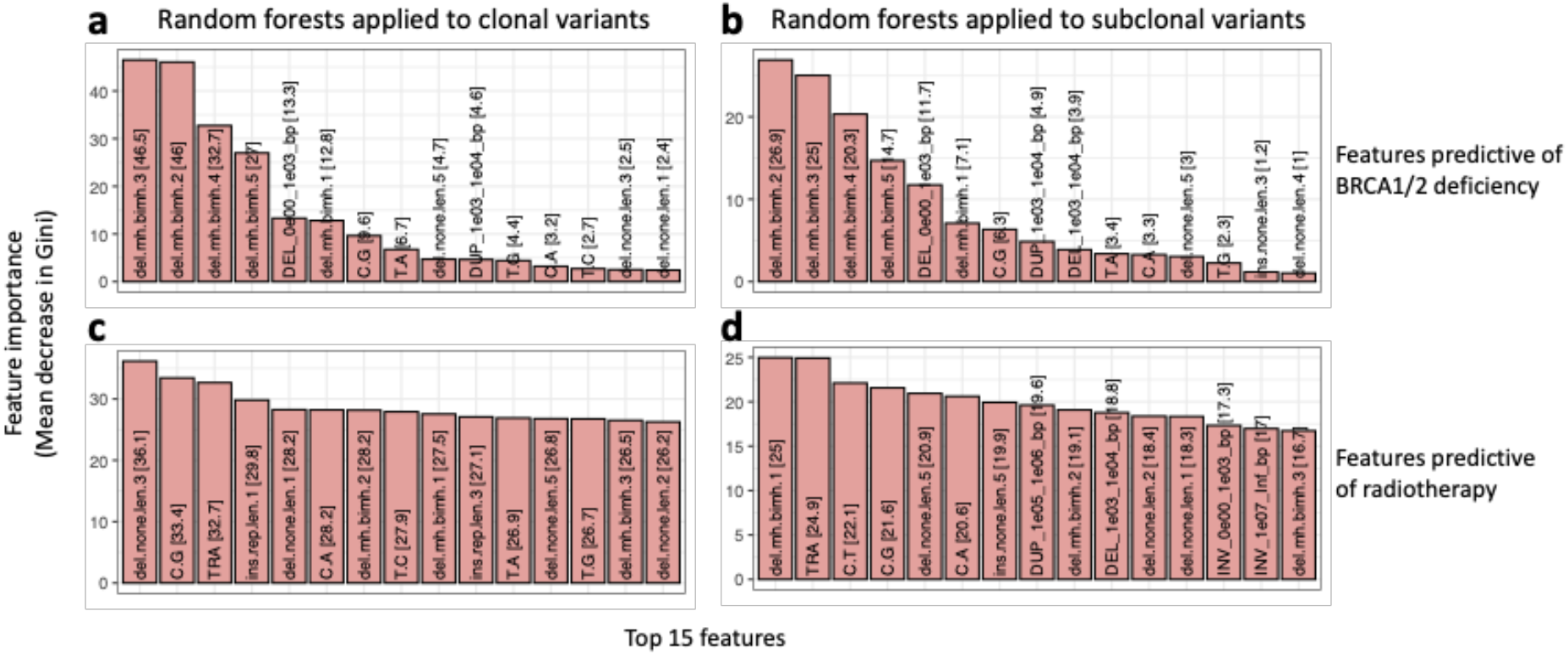
Random forests were used to identify the features predictive of BRCA1/2 deficiency (**a,b**) compared to those predictive of radiotherapy (**c,d**) when using clonal versus subclonal variants as input. Feature importance is indicated by mean decrease in Gini, with only the top 15 important features being shown. Deletions with 1bp of flanking homology (del.mh.bimh.1) was more associated with radiotherapy especially in the subclonal fraction, while deletions with ≥2bp flanking homology (del.mh.bimh.2.5) was more associated with BRCA1/2 deficiency.

**Supplementary figure 13:**
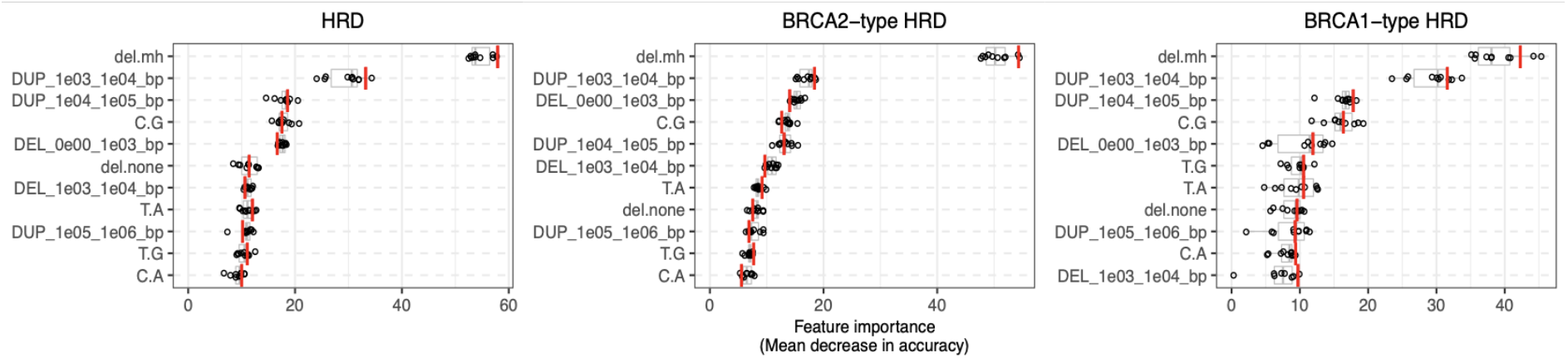
The features used by CHORD-del.mh.merged to predict HRD as well as BRCA1-type HRD and BRCA2-type HRD, with their importance indicated by mean decrease in accuracy. Deletions with flanking microhomology (del.mh) was the most important feature for predicting HRD as a whole, with 1-100kb structural duplications (DUP_1e03_1e04_bp, DUP_1e04_1e05_bp) differentiating BRCA1-type HRD from BRCA2-type HRD. Boxplot and dots show the feature importance over 10-folds of nested CV on the training set, with the red line showing the feature importance in the final CHORD model. Boxes show the interquartile range (IQR) and whiskers show the largest/smallest values within 1.5 times the IQR.

**Supplementary figure 14:**
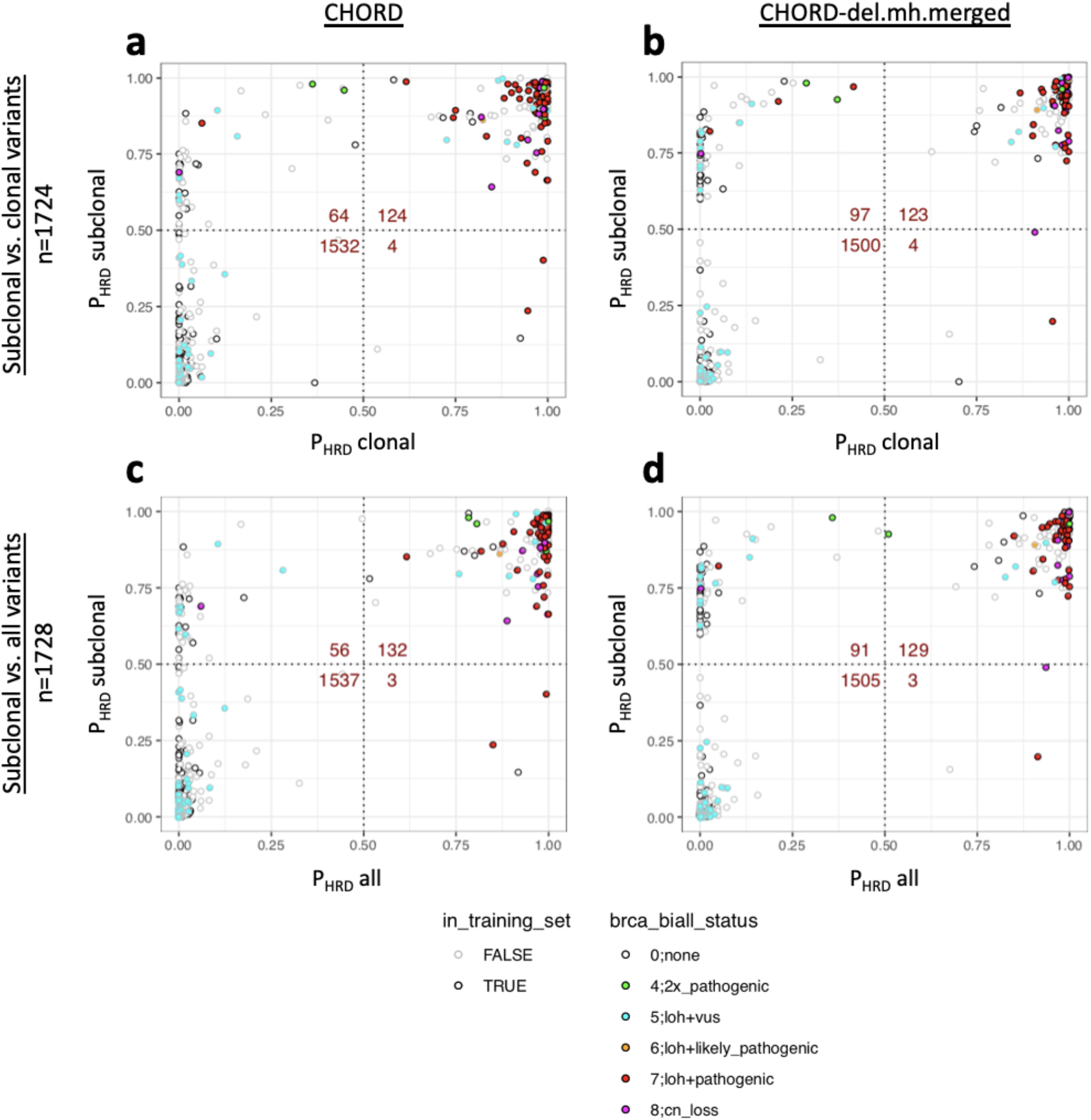
HRD predictions (on HMF samples) on (**a,b**) subclonal versus clonal variants and (**c,d**) subclonal vs. all variants from CHORD and CHORD-del.mh.merged. Only samples passing CHORD’s QC criteria were shown (MSI negative, ≥50 indels in both clonal and subclonal fractions, and ≥30 SVs if a sample was predicted HRD). Dots with a black outline belonged to the training set of CHORD. Dot fill color indicates the biallelic status of BRCA1 or BRCA2.

**Supplementary figure 15:**
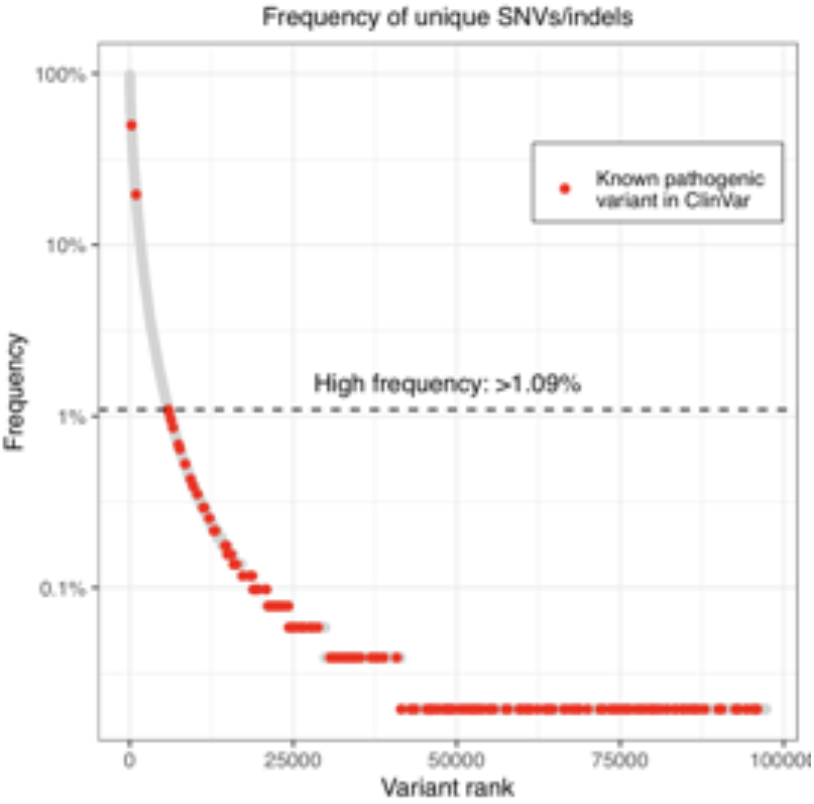
Frequency of unique germline SNVs/indels of 781 cancer related genes in patients of the HMF cohort. Germline variants with a frequency >1.09% were marked as benign prior to performing the pan-cancer analysis of HRD. Two known pathogenic variants above this frequency were also considered benign.

**Supplementary figure 16:**
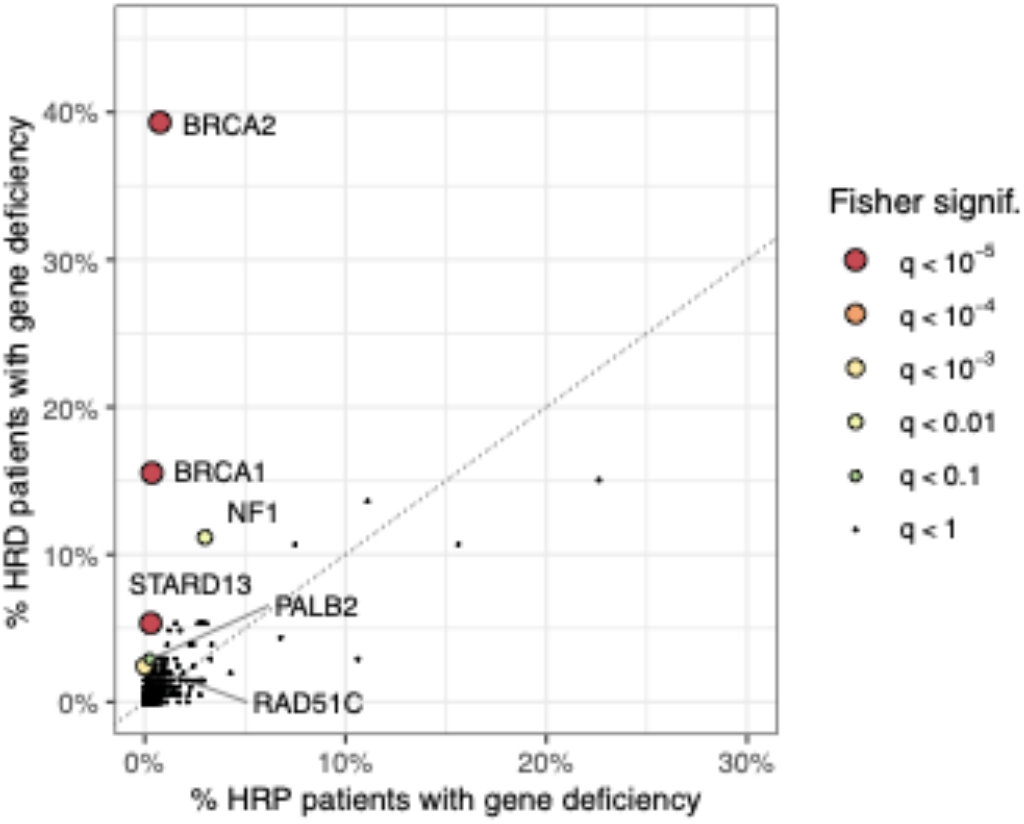
Same as **Figure 3b** except with NF1 and STARD13 included. A one-tailed Fisher’s exact test determined BRCA1, BRCA2, RAD51C, PALB2, NF1 and STARD13 to be significantly enriched (from a list of 781 cancer related genes) in CHORD-HRD vs. CHORD-HRP patient groups. Each point represents a gene with its size/color corresponding to the statistical significance as determined by the Fisher’s exact test, with axes indicating the percentage of patients (within either the CHORD-HRD or CHORD-HRP group) in which biallelic inactivation was detected. Multiple testing correction was performed using the Hochberg procedure.

**Supplementary figure 17:**
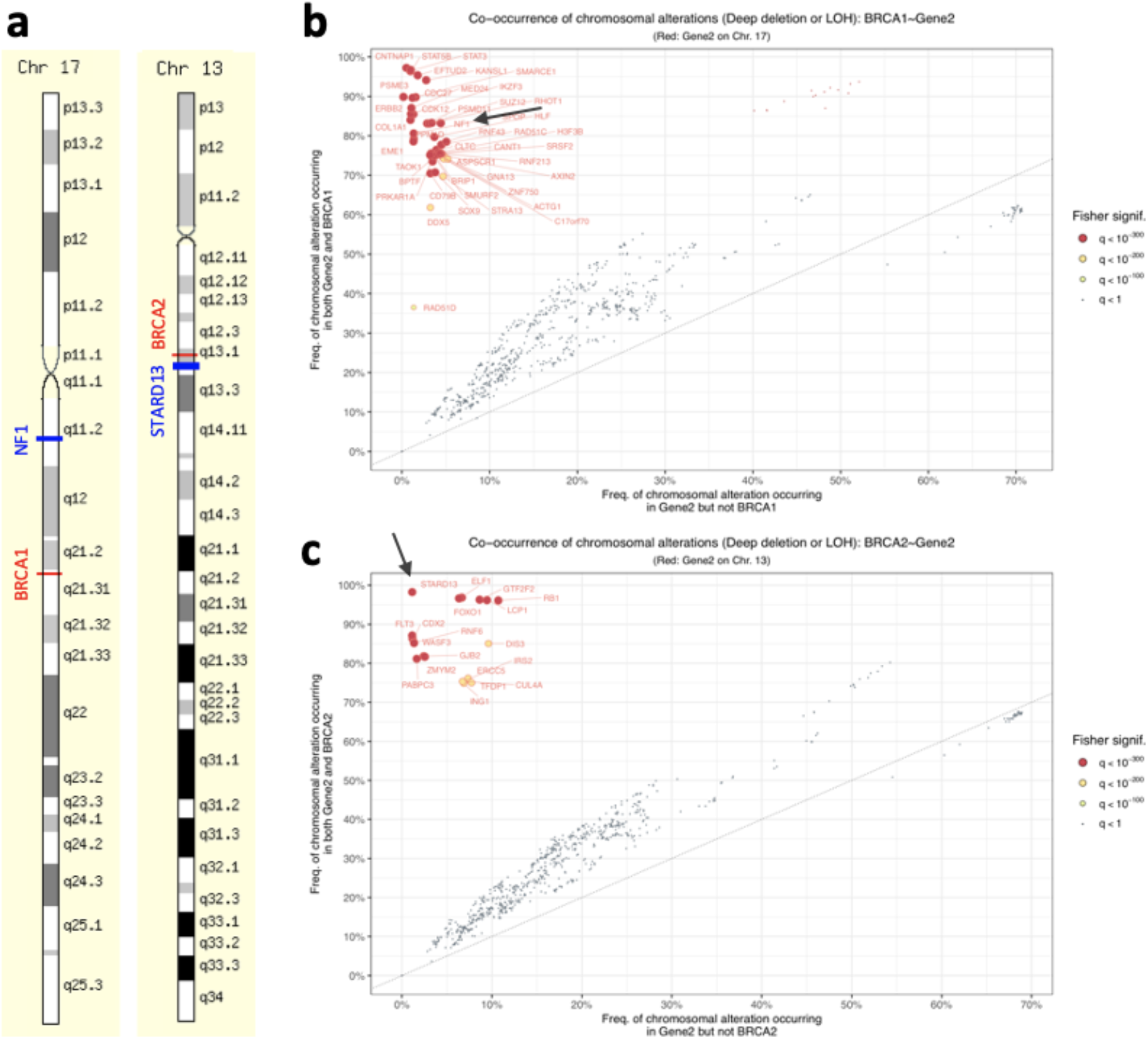
Chromosomal alterations (deep deletions or loss of heterozygosity (LOH)) affecting BRCA1 and BRCA2 also affects nearby genes including NF1 and STARD13 respectively. (**a**) BRCA1 and NF1 are both located on Chr17q while BRCA2 and STARD13 are both located on Chr13q. Source: www.genecards.org. (**b**) Enrichment of NF1 biallelic loss as shown in **Supplementary figure 16** is likely due to the gene being within proximity of BRCA1 and not because the gene is associated with HRD, since NF1 is not considered to be involved in HR in literature. The size/color of each point on the plot represents the significance of enrichment of CNAs occurring both in BRCA1 and in each of the 781 genes (one vs. all comparison). This was determined by a one-tailed Fisher’s exact test, where multiple testing correction was performed using the Hochberg procedure. Genes residing on the same chromosome as BRCA1 were marked with a red outline and text. (**c**) Similarly, as with (**a**), a chromosomal alteration affecting BRCA2 also affects the nearby gene STARD13. Enrichment of STARD13 biallelic loss as shown in **Supplementary figure 16** is likely due to the gene being within proximity of BRCA2 and not because the gene is associated with HRD.

**Supplementary figure 18:**
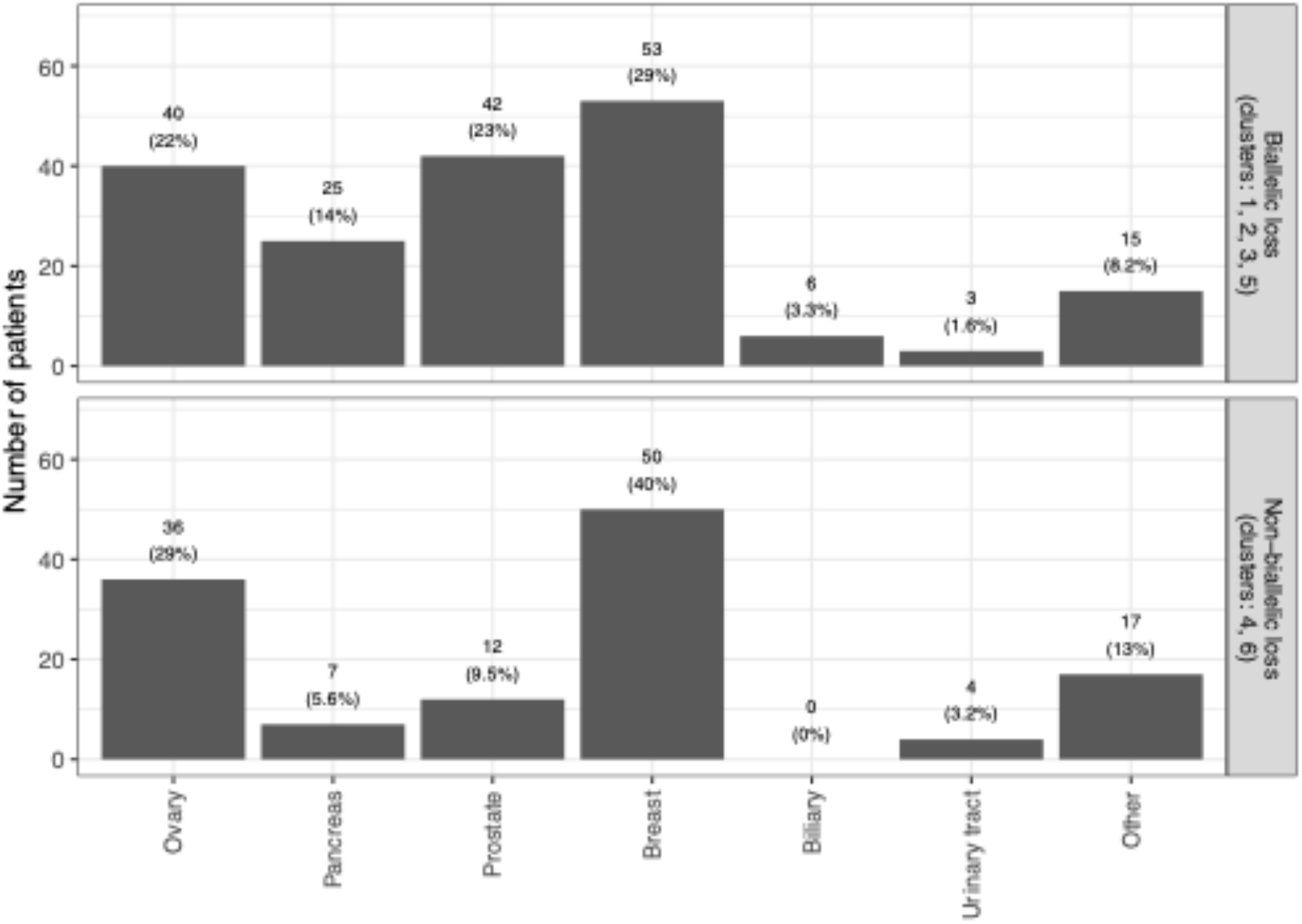
The number of CHORD-HRD patients of each cancer type which did (top) and did not (bottom) have biallelic loss of BRCA1, BRCA2, RAD51C, or PALB2. The top panel corresponds to patients in cluster 1,2,3 and 5 of **Figure 3c** while the bottom panel corresponds to patients in cluster 4 and 6. The percentages shown are within-group proportions.

**Supplementary figure 19:**
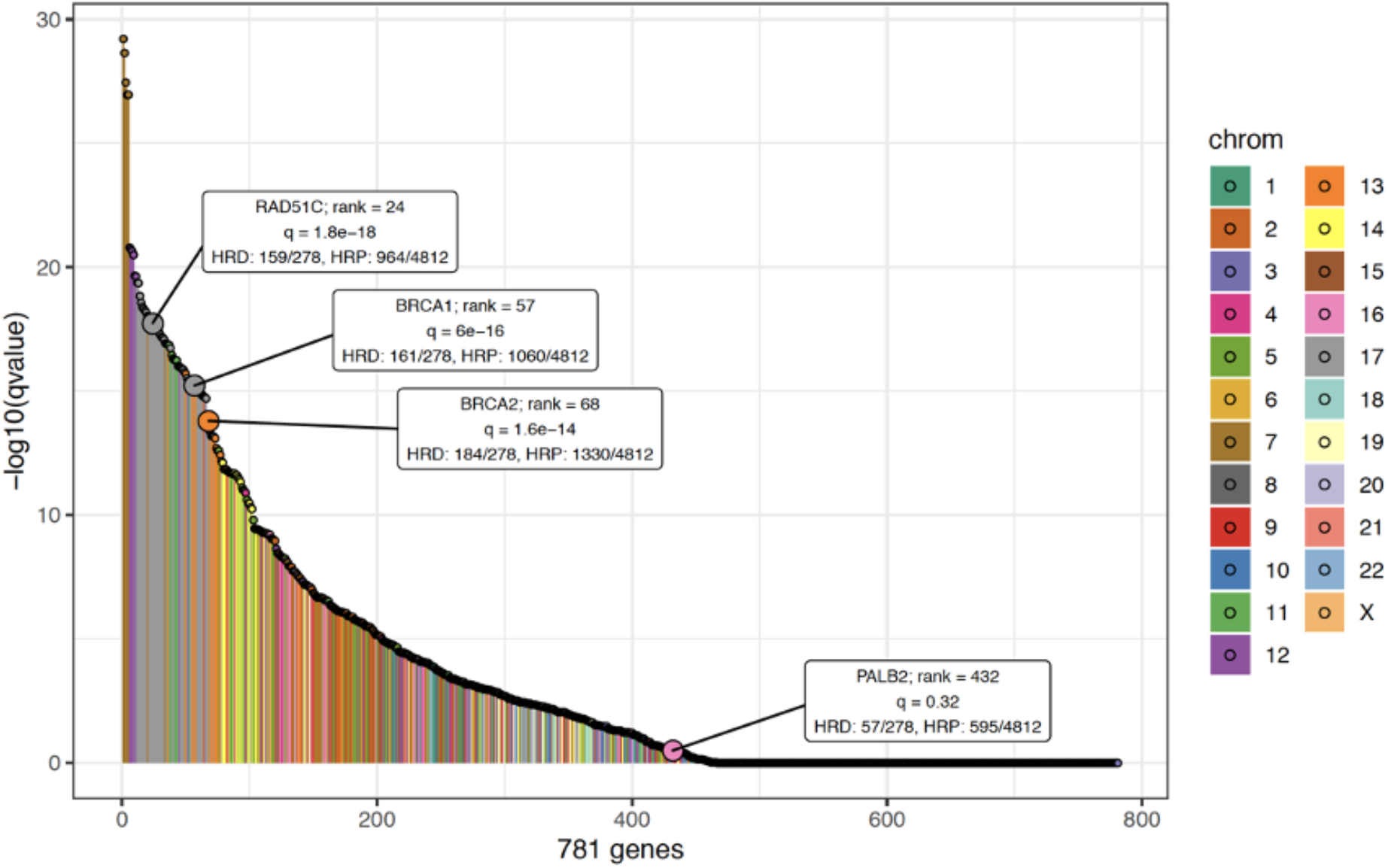
Enrichment of LOH in the 781 HR/cancer related genes between CHORD-HRD samples (excluding those with deep deletions in BRCA1/2, RAD51C or PALB2) and CHORD-HRP samples. Enrichment was calculated for each gene using a one-sided Fisher’s exact test.

**Supplementary figure 20:**
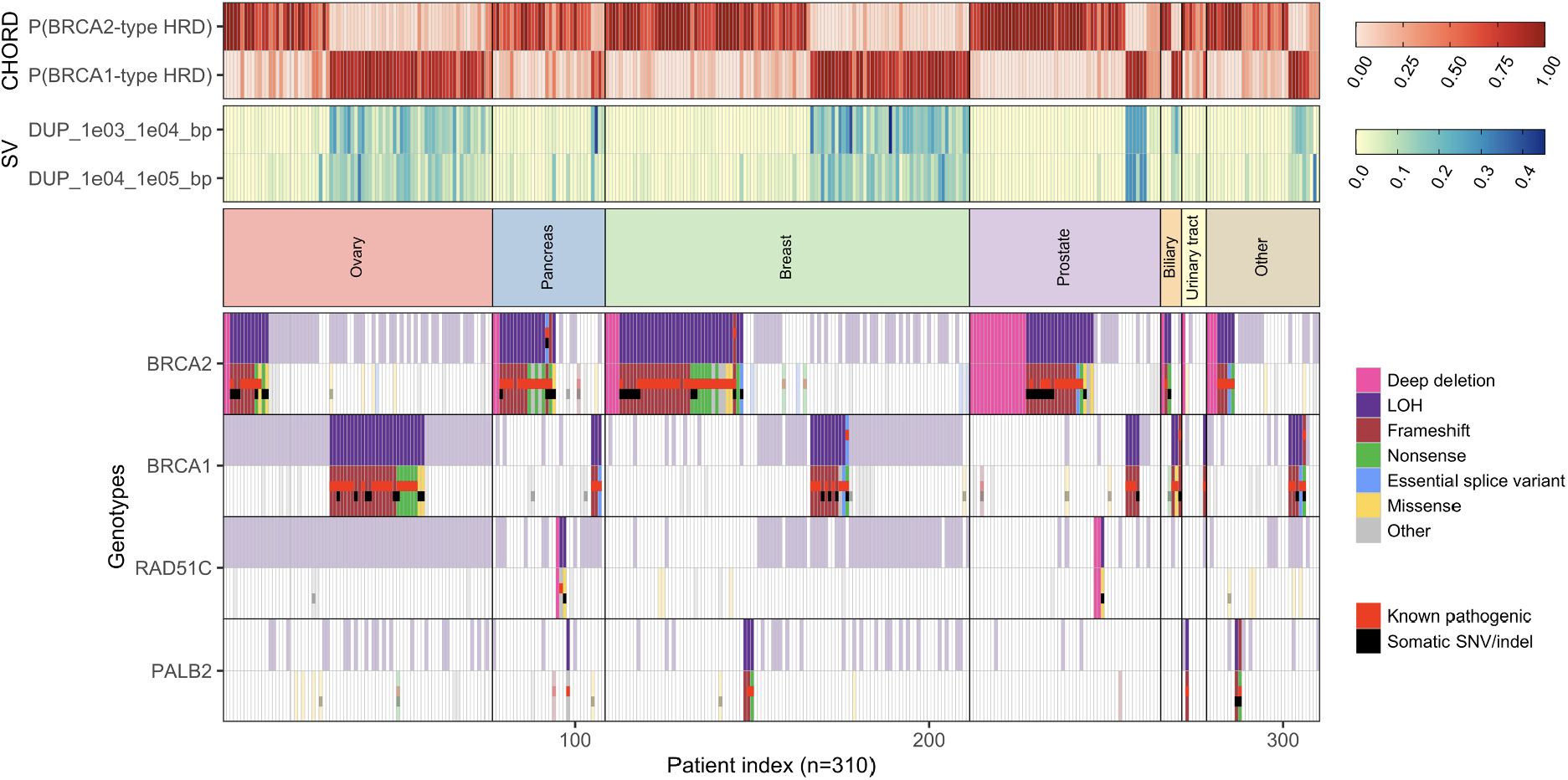
Biallelic status of BRCA2, BRCA1, RAD51C and PALB2 in CHORD-HRD patients from both the HMF and PCAWG datasets. Patients were clustered both by cancer type and by HRD type. Top: BRCA1- and BRCA2-type HRD probabilities from CHORD. Middle: SV contexts used by CHORD to distinguish BRCA1- from BRCA2-type HRD. Bottom: The biallelic status of each gene. Tiles marked as ‘Known pathogenic’ refer to variants having a ‘pathogenic’ or ‘likely pathogenic’ annotation in ClinVar. Only data from samples that passed CHORD’s QC criteria are shown in this figure (MSI absent, ≥50 indels, and ≥30 SVs if a sample was predicted HRD).

**Supplementary figure 21:**
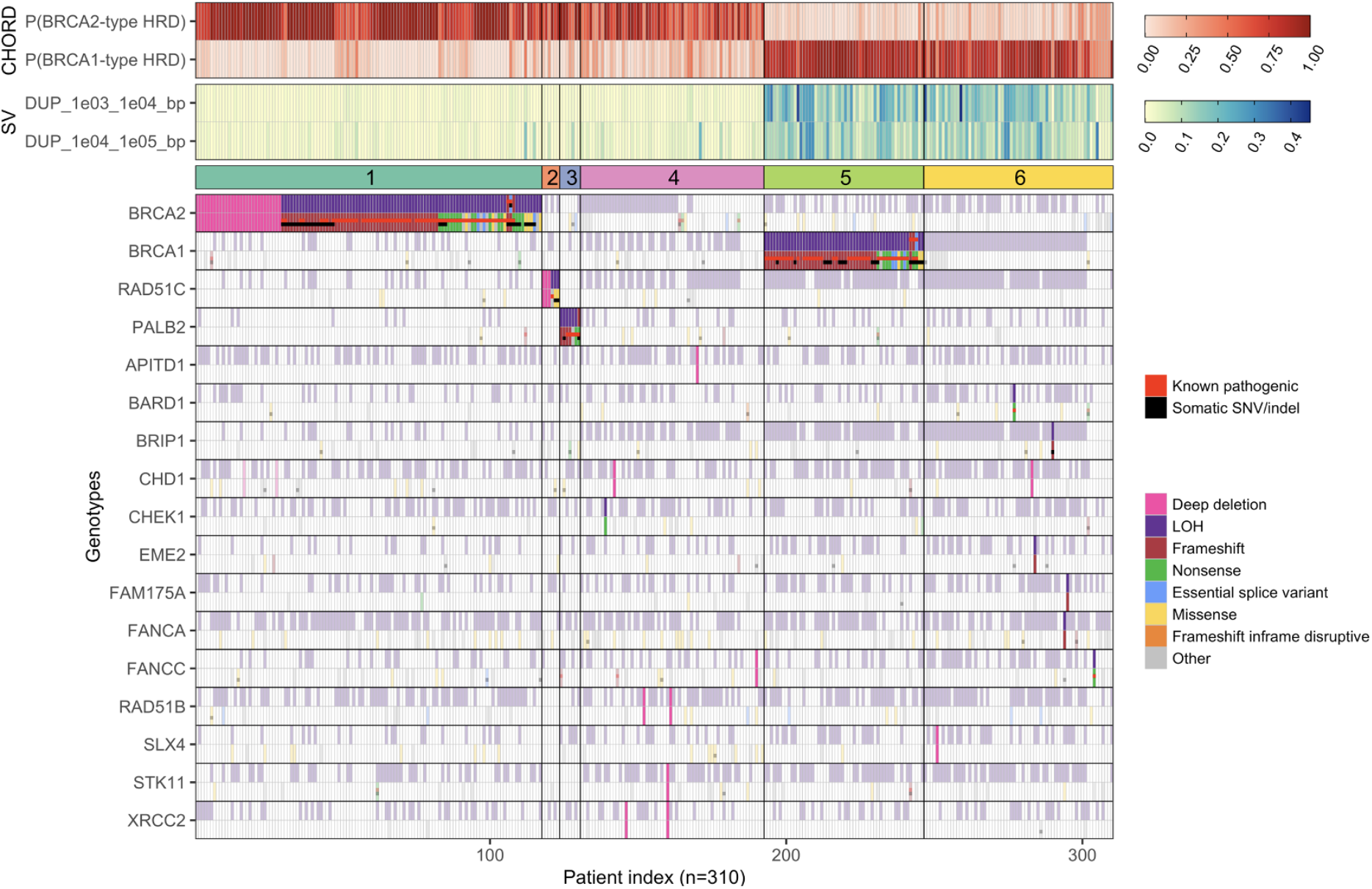
Biallelic status of BRCA2, BRCA1, RAD51C, PALB2, as well as other HR genes in CHORD-HRD patients from both the HMF and PCAWG datasets. Top: BRCA1- and BRCA2-type HRD probabilities from CHORD. Middle: SV contexts used by CHORD to distinguish BRCA1- from BRCA2-type HRD. Bottom: The biallelic status of each gene. Tiles marked as ‘Known pathogenic’ refer to variants having a ‘pathogenic’ or ‘likely pathogenic’ annotation in ClinVar. Only HR genes with one of the following events in at least one patient from cluster 4 or 6 was shown here: deep deletion; or LOH in combination with a pathogenic/likely pathogenic variant or a frameshift/nonsense variant. Only data from samples that passed CHORD’s QC criteria are shown in this figure (MSI absent, ≥50 indels, and ≥30 SVs if a sample was predicted HRD). Furthermore, only genes where at least one patient had an impactful biallelic event are shown.

**Supplementary figure 22:**
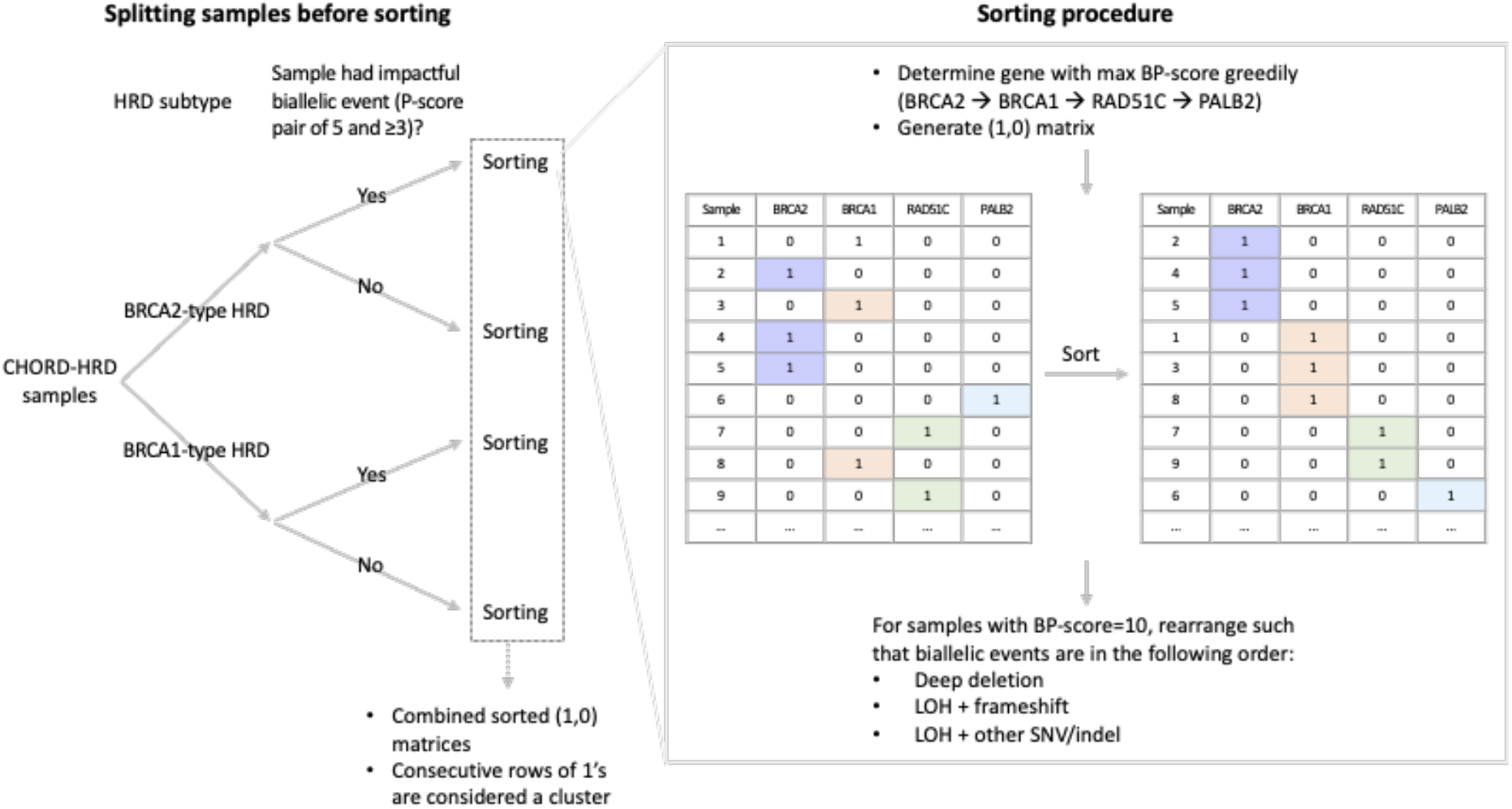
Overview of the procedure to sort and cluster CHORD-HRD samples.

## References

1. Lord CJ, Ashworth A. BRCAness revisited. Nat Rev Cancer. 2016;16:110–20.

2. Heeke A, Lynce F, Baker T, Pishvaian M, Isaacs C. Prevalence of Homologous Recombination Deficiency (HRD) Among All Tumor Types. 2017.

3. Audeh MW, Carmichael J, Penson RT, Friedlander M, Powell B, Bell-McGuinn KM, et al. Oral poly(ADP-ribose) polymerase inhibitor olaparib in patients with BRCA1 or BRCA2 mutations and recurrent ovarian cancer: a proof-of-concept trial. The Lancet. 2010;376:245–51.

4. Mateo J, Carreira S, Sandhu S, Miranda S, Mossop H, Perez-Lopez R, et al. DNA-Repair Defects and Olaparib in Metastatic Prostate Cancer.http://dx.doi.org/101056/NEJMoa1506859. 2015. doi:10.1056/NEJMoa1506859.

5. Hoppe MM, Sundar R, Tan DSP, Jeyasekharan AD. Biomarkers for Homologous Recombination Deficiency in Cancer. JNCI J Natl Cancer Inst. 2018;110:704–13.

6. Van Hoeck A, Tjoonk NH, van Boxtel R, Cuppen E. Portrait of a cancer: mutational signature analyses for cancer diagnostics. BMC Cancer. 2019;19:457.

7. Sfeir A, Symington LS. Microhomology-mediated end joining: a back-up survival mechanism or dedicated pathway? Trends Biochem Sci. 2015;40:701–14.

8. Nik-Zainal S, Davies H, Staaf J, Ramakrishna M, Glodzik D, Zou X, et al. Landscape of somatic mutations in 560 breast cancer whole-genome sequences. Nature. 2016;534:47–54.

9. Davies H, Glodzik D, Morganella S, Yates LR, Staaf J, Zou X, et al. HRDetect is a predictor of *BRCA1* and *BRCA2* deficiency based on mutational signatures. Nat Med. 2017;23:517–25.

10. Degasperi A, Amarante TD, Czarnecki J, Shooter S, Zou X, Glodzik D, et al. A practical framework and online tool for mutational signature analyses show intertissue variation and driver dependencies. Nat Cancer. 2020;1:249–63.

11. Nones K, Johnson J, Newell F, Patch AM, Thorne H, Kazakoff SH, et al. Whole-genome sequencing reveals clinically relevant insights into the aetiology of familial breast cancers. Ann Oncol. 2019;30:1071–9.

12. Staaf J, Glodzik D, Bosch A, Vallon-Christersson J, Reuterswärd C, Häkkinen J, et al. Whole-genome sequencing of triple-negative breast cancers in a population-based clinical study. Nat Med. 2019;25:1526–33.

13. Priestley P, Baber J, Lolkema MP, Steeghs N, Bruijn E de, Shale C, et al. Pan-cancer whole-genome analyses of metastatic solid tumours. Nature. 2019;:1–7.

14. Campbell PJ, Getz G, Korbel JO, Stuart JM, Jennings JL, Stein LD, et al. Pan-cancer analysis of whole genomes. Nature. 2020;578:82–93.

15. Jonsson P, Bandlamudi C, Cheng ML, Srinivasan P, Chavan SS, Friedman ND, et al. Tumour lineage shapes BRCA-mediated phenotypes. Nature. 2019;571:576–9.

16. Polak P, Kim J, Braunstein LZ, Karlic R, Haradhavala NJ, Tiao G, et al. A mutational signature reveals alterations underlying deficient homologous recombination repair in breast cancer. Nat Genet. 2017;49:1476–86.

17. Gulhan DC, Lee JJ-K, Melloni GEM, Cortés-Ciriano I, Park PJ. Detecting the mutational signature of homologous recombination deficiency in clinical samples. Nat Genet. 2019;51:912–9.

18. Maura F, Degasperi A, Nadeu F, Leongamornlert D, Davies H, Moore L, et al. A practical guide for mutational signature analysis in hematological malignancies. Nat Commun. 2019;10:1–12.

19. Pich O, Muiños F, Lolkema MP, Steeghs N, Gonzalez-Perez A, Lopez-Bigas N. The mutational footprints of cancer therapies. Nat Genet. 2019;51:1732–40.

20. Kucab JE, Zou X, Morganella S, Joel M, Nanda AS, Nagy E, et al. A Compendium of Mutational Signatures of Environmental Agents. Cell. 2019;177:821–836.e16.

21. Behjati S, Gundem G, Wedge DC, Roberts ND, Tarpey PS, Cooke SL, et al. Mutational signatures of ionizing radiation in second malignancies. Nat Commun. 2016;7:1–8.

22. Christensen S, Van der Roest B, Besselink N, Janssen R, Boymans S, Martens JWM, et al. 5-Fluorouracil treatment induces characteristic T>G mutations in human cancer. Nat Commun. 2019;10:4571.

23. Póti Á, Gyergyák H, Németh E, Rusz O, Tóth S, Kovácsházi C, et al. Correlation of homologous recombination deficiency induced mutational signatures with sensitivity to PARP inhibitors and cytotoxic agents. Genome Biol. 2019;20:240.

24. Chun J, Buechelmaier ES, Powell SN. Rad51 Paralog Complexes BCDX2 and CX3 Act at Different Stages in the BRCA1-BRCA2-Dependent Homologous Recombination Pathway. Mol Cell Biol. 2013;33:387–95.

25. Zhao W, Steinfeld JB, Liang F, Chen X, Maranon DG, Jian Ma C, et al. BRCA1–BARD1 promotes RAD51-mediated homologous DNA pairing. Nature. 2017;550:360–5.

26. Cantor SB, Guillemette S. Hereditary breast cancer and the BRCA1-associated FANCJ/BACH1/BRIP1. Future Oncol Lond Engl. 2011;7:253–61.

27. Castillo A, Paul A, Sun B, Huang TH, Wang Y, Yazinski SA, et al. The BRCA1-Interacting Protein Abraxas Is Required for Genomic Stability and Tumor Suppression. Cell Rep. 2014;8:807–17.

28. Folias A, Matkovic M, Bruun D, Reid S, Hejna J, Grompe M, et al. BRCA1 interacts directly with the Fanconi anemia protein FANCA. Hum Mol Genet. 2002;11:2591–7.

29. Quigley DA, Dang HX, Zhao SG, Lloyd P, Aggarwal R, Alumkal JJ, et al. Genomic Hallmarks and Structural Variation in Metastatic Prostate Cancer. Cell. 2018;174:758–769.e9.

30. Golan T, Hammel P, Reni M, Van Cutsem E, Macarulla T, Hall MJ, et al. Maintenance Olaparib for Germline BRCA-Mutated Metastatic Pancreatic Cancer. N Engl J Med. 2019;381:317–27.

31. Pilarski R. The Role of BRCA Testing in Hereditary Pancreatic and Prostate Cancer Families. Am Soc Clin Oncol Educ Book. 2019;:79–86.

32. Chopra N, Tovey H, Pearson A, Cutts R, Toms C, Proszek P, et al. Homologous recombination DNA repair deficiency and PARP inhibition activity in primary triple negative breast cancer. Nat Commun. 2020;11:1–12.

33. Arora K, Barbieri C. Molecular Subtypes of Prostate Cancer. Curr Oncol Rep. 2018;20. doi:https://doi.org/10.1007/s11912-018-0707-9.

34. Tariq N-A, McNamara M, Valle J. Biliary tract cancers: current knowledge, clinical candidates and future challenges. Cancer Manag Res. 2019;11:2623–2642.

35. Sakai W, Swisher EM, Karlan BY, Agarwal MK, Higgins J, Friedman C, et al. Secondary mutations as a mechanism of cisplatin resistance in BRCA2-mutated cancers. Nature. 2008;451:1116–20.

36. Edwards SL, Brough R, Lord CJ, Natrajan R, Vatcheva R, Levine DA, et al. Resistance to therapy caused by intragenic deletion in BRCA2. Nature. 2008;451:1111–5.

37. Nangalia J, Campbell PJ. Genome Sequencing during a Patient’s Journey through Cancer. N Engl J Med. 2019;381:2145–56.

38. Cameron DL, Baber J, Shale C, Papenfuss AT, Valle-Inclan JE, Besselink N, et al. GRIDSS, PURPLE, LINX: Unscrambling the tumor genome via integrated analysis of structural variation and copy number. bioRxiv. 2019;:781013.

